# Measurement of Volatile Compounds for Real-time Analysis of Soil Microbial Metabolic Response to Simulated Snowmelt

**DOI:** 10.1101/2021.03.11.432778

**Authors:** Junhyeong Kim, Allen H. Goldstein, Romy Chakraborty, Kolby Jardine, Robert Weber, Patrick O. Sorensen, Shi Wang, Boris Faybishenko, Pawel K. Misztal, Eoin L. Brodie

**Affiliations:** Lawrence Berkeley National Laboratory, Climate and Ecosystems Sciences, Earth and Environmental Sciences, Berkeley, CA, USA; Department of Environmental Science, Policy and Management, University of California Berkeley, Berkeley, CA, USA; Department of Civil, Architectural and Environmental Engineering, University of Texas Austin, Austin, TX, USA

**Author notes:** **Correspondence:** Eoin L. Brodie.

## Abstract

Snowmelt dynamics are a significant determinant of microbial metabolism in soil and regulate global biogeochemical cycles of carbon and nutrients by creating seasonal variations in soil redox and nutrient pools. With an increasing concern that climate change accelerates both snowmelt timing and rate, obtaining an accurate characterization of microbial response to snowmelt is important for understanding biogeochemical cycles intertwined with soil. However, observing microbial metabolism and its dynamics non-destructively remains a major challenge for systems such as soil. Microbial volatile compounds (mVCs) emitted from soil represent information-dense signatures and when assayed non-destructively using state-of-the-art instrumentation such as Proton Transfer Reaction-Time of Flight-Mass Spectrometry (PTR-TOF-MS) provide time resolved insights into the metabolism of active microbiomes. In this study, we used PTR-TOF-MS to investigate the metabolic trajectory of microbiomes from a subalpine forest soil, and their response to a simulated wet-up event akin to snowmelt. Using an information theory approach based on the partitioning of mutual information, we identified mVC metabolite pairs with robust interactions, including those that were non-linear and with time lags. The biological context for these mVC interactions was evaluated by projecting the connections onto the Kyoto Encyclopedia of Genes and Genomes (KEGG) network of known metabolic pathways. Simulated snowmelt resulted in a rapid increase in the production of trimethylamine (TMA) suggesting that anaerobic degradation of quaternary amine osmo/cryoprotectants, such as glycine betaine, may be important contributors to this resource pulse. Unique and synergistic connections between intermediates of methylotrophic pathways such as dimethylamine, formaldehyde and methanol were observed upon wet-up and indicate that the initial pulse of TMA was likely transformed into these intermediates by methylotrophs. Increases in ammonia oxidation signatures (transformation of hydroxylamine to nitrite) were observed in parallel, and while the relative role of nitrifiers or methylotrophs cannot be confirmed, the inferred connection to TMA oxidation suggests either a direct or indirect coupling between these processes. Overall, it appears that such mVC time-series from PTR-TOF-MS combined with causal inference represents an attractive approach to non-destructively observe soil microbial metabolism and its response to environmental perturbation.

## 1 Introduction

Soil is a major compartment of the Earth system and its constituent organisms are central contributors to global biogeochemical cycles. Accurate characterization of the activity of soil microbiomes is key to understanding how the soil compartment functions and responds to chronic and acute perturbations (Delgado-Baquerizo et al., 2019; Bardgett et al., 2014). Snowpack dynamics for example are strongly influenced by climate driving forces such as atmospheric warming and precipitation patterns, and in turn regulate both plant and microbial activity (Wipf, 2010; Steltzer et al., 2009; Brooks et al., 1998; Lipson et al., 2002; Sorensen et al., 2019). Our prior work observed that snowmelt induced a significant shift in soil water potential, dynamic biomass growth and turnover that corresponded to a reassembly of the soil microbiome (Sorensen et al., 2020). During this critical phase of the water year in snow dominated systems, nitrogen retention and release by microbial biomass is dynamic and associated with snowmelt dynamics (Brooks et al., 1998). This response to snowmelt infiltration is similar to the Birch Effect that has been observed as the respiration response of a dry soil to wet-up (Birch, 1958). This response is rapid with turnover of microbial biomass, community composition, gene expression and metabolism being highly dynamic over a period of a few hours to days (e.g. Placella et al., 2012; Placella et al., 2013; Blazewicz et al., 2020; Barnard et al., 2020). Capturing such microbial metabolic trajectories in heterogeneous systems like soils remains challenging due to problems associated with destructive sampling and low time resolution.

Although the characterization of soil microbial metabolism has become much more convenient and effective due to the development of sequence-based and mass spectrometry approaches to interrogate biomolecular processes, most approaches require destructive sample retrieval and processing. This limits analyses to sequential snapshots rather than more continuous time-series and results in important information loss. Non-destructive approaches such as the continuous monitoring of trace gases can be considered ‘information-rich,’ although not ‘information-dense,’ and therefore provide more limited insight into mechanistic processes underlying their fluxes. While sequence-based or metabolomics approaches requiring extraction excel at delivering broad snapshots of soil microbiome function, alternative and non-destructive approaches to monitor microbial metabolic dynamics are needed.

Microbial volatile compounds (mVCs) represent a range of organic and inorganic compounds that can exist in both aqueous and gaseous phases under standard conditions. Together with recent advances in measurement techniques, mVCs show great promise in non-destructive representation of microbial metabolism across multiple disciplines, including soil microbiology and epidemiology (McNeal and Herbert, 2009; Palma et al., 2018). mVCs can diffuse through cell membranes and circulate through and from soil (Stotzky et al. 1976). The high mobility of soil mVCs allows them to act as substrates and inhibitors of microbial metabolism (Tyc et al., 2017; Netzker et al., 2020) as well as signalling molecules over large distances (Schulz-Bohm and Martin-Sanchez, 2017). Proton transfer reaction time of flight mass spectrometer (PTR-TOF-MS) is a state-of-the-art instrument for real-time mVC measurement and can be used to quantify mVC composition and dynamics over timescales of seconds to days, providing the opportunity to observe these moderately information-dense signatures of microbial metabolism in real-time (Seewald et al., 2010).

This combination of information-rich time series and information-dense measurements may provide opportunities to explore mVC interactions in a metabolic network context. These metabolic networks are dynamic in terms of interaction strength and connectivity including both linear and non-linear relationships (Goodwell and Kumar 2017). Approaches based on information theory have shown promise to identify robust and potentially causal interactions in applications with similar data properties (Goodwell and Kumar., 2017; Jiang and Kumar, 2019; Goodwell et al., 2020).

In this study, we evaluate the potential of using mVCs to follow the dynamics of microbial metabolic pathways in microcosm experiments where soils from a subalpine conifer forest in Colorado, USA were subjected to a wet-up akin to snowmelt. First, we measured mVC profiles of microcosm headspace in a continuous flow system via PTR-TOF-MS and used these mVC time-series to establish directed connections between various mVCs. Using an information theory approach, we inferred the directed connections as causal (Goodwell and Kumar, 2017). Subsequently, we created a reaction network of mVCs mapped onto the existing KEGG (Kyoto Encyclopedia of Genes and Genomes) reaction network to explore the biological relevance of observed connections based on prior knowledge (Kanehisa, 2000). The observed metabolic connections are interpreted in the context of snowmelt and its impact on soil microbial metabolism. We expected to see the emergence of biological pathways in VC-based molecular networks upon wet-up. These pathways would include degradation of microbial metabolites that are closely related to osmotic regulation, as well as litter-derived substrates that become incorporated into central metabolism. Biological reactions with multiple intermediates were hypothesized to display not only strong unique (U), but also synergistic (S) information to more accurately reflect the complexity of observed metabolic pathways.

## 2 Materials and Methods

### Soil properties and microcosm preparation

Soils were sampled on February 19th 2019 from a Spruce pine dominated hillslope located at Snodgrass Mountain, CO (38°55’55.60″N 106°59’9.82″W) in parallel with snow excavation for snow water equivalent analyses. Snow depth was 163cm at the time of sampling and soil cores were taken to a depth of 15 cm from the soil surface. Dissolved inorganic nitrogen pools were determined colorimetrically according to protocols as described previously (Sims et al., 1995; Horwath et al. 2003). NO_3_^−^ concentration was below detection limit and NH_4_^+^ concentration was 1.91 ± 0.07 μM. Soil water content was measured gravimetrically at 0.136 g H_2_O/g soil. A water retention curve was obtained and used to determine soil water potential, estimated at −1.5 MPa for these samples prior to wet-up. The retention curve was measured with an HYPROP2 by METER. HYPROP2 uses the evaporation method to investigate the relationship between water content and the associated matric potential from 0 kPa to −100 kPa (Schindler et al., 2010a,b; Peters et al., 2015). This matric potential range covers full saturation condition, field capacity, and wilting point, providing useful information on the actual water availability in soil. The van Genuchten-Mualem model was used to approximate soil hydraulic properties from retention curve measurement (van Genuchten, 1980; van Genuchten and Nielsen, 1985). Before mVC measurement, the soil microcosms were left at room temperature to acclimate for 12 hours and the headspace of each microcosm was flushed under 70 mL/min of VC-free air (zero-air) for 1 hour.

In order to investigate the effects of wet-up on the soil mVC profile, soil microcosms were prepared in 300 mL VC-grade glass jars with Teflon covers. Each jar contained 20 g soil wet weight with an empty jar used to control for any background VC fluxes (Fig. 1). Microcosms were initially maintained at 80% relative humidity to simulate soil underlying snow cover for 6 days to establish mVC profiles prior to a simulated snowmelt wet-up event. For soil wet-up, milli-Q water was sterilized by membrane filtration (0.2 μm pore size) and 9.0 mL added to each jar to achieve approximately 75% soil water saturation. Measurement of mVC fluxes continued for an additional 3 days following this wet-up event.

**Figure 1.**
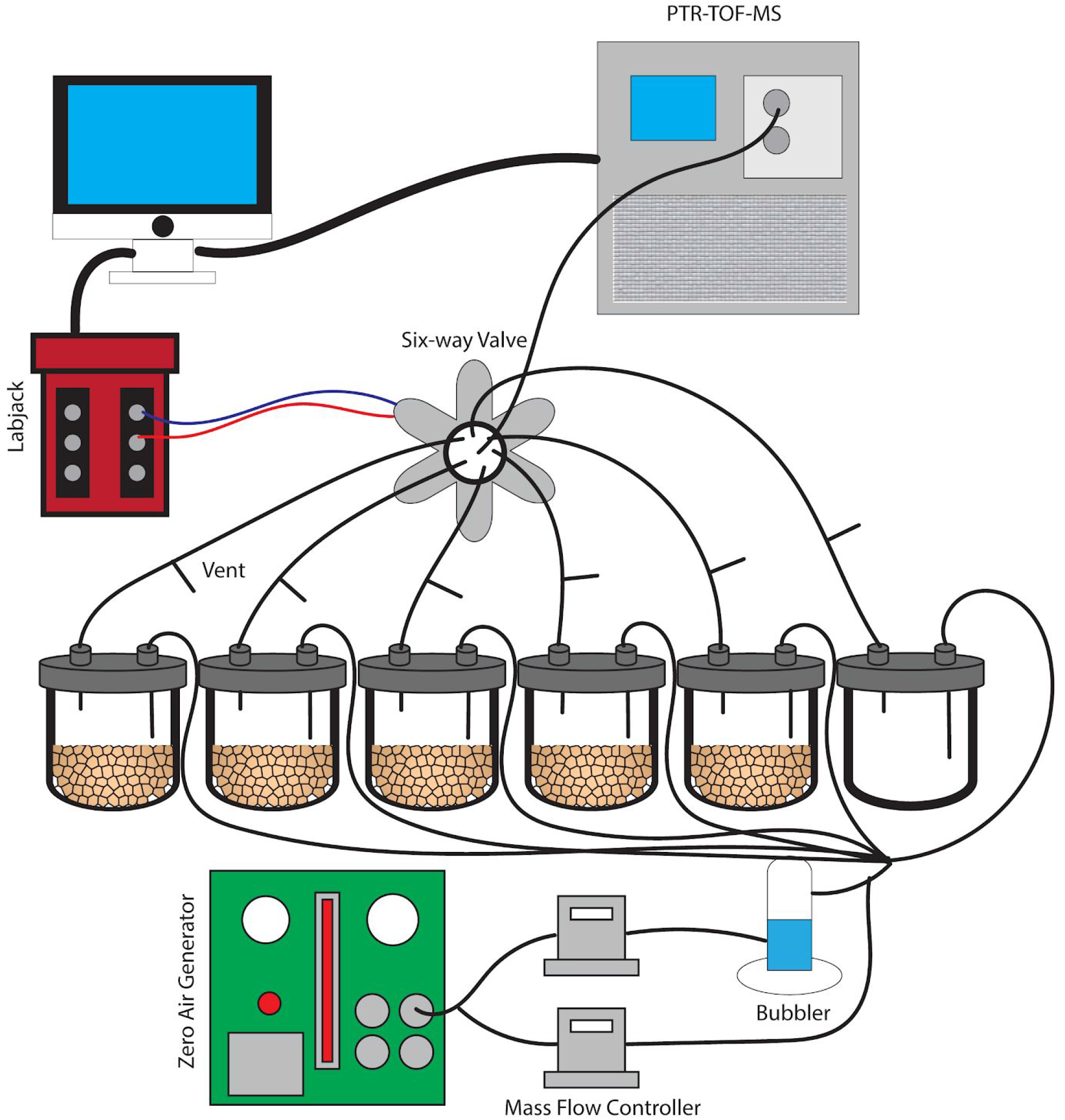
A schematic image of the dynamic flow-through system used for soil VC measurement. Soils were incubated without water for 6 days and were subsequently saturated with water. This set-up provides clean and humidified air to the soils and also prevents artificial accumulation of trace gases with constant outflow matching the inflow air.

### Microcosm Flow-through System

A dynamic flow-through gas-exchange system was used to maintain an 80% humidity following wetting and avoid unnatural accumulation of mVCs and trace gases such as CO_2_ during microcosm incubations (Misztal et al., 2018). A constant inflow of air into each of the 6 jars was matched by an equal outflow. The airflow did not pass through the soil itself, but rather mixed with the soil headspace air. At the outlet of each jar, a vent was installed to maintain atmospheric pressure conditions, continuous airflow, and to mimic the diffusion of mVCs from the soil into a mixed atmosphere. Five soil replicates plus an empty jar for a control were installed in parallel, each with a constant flow velocity of 70mL/min (Fig. 1). The inflow air was purified of organic compounds using an Aadco 737 Zero Air Generator (Aadco Inst, Cleves, OH) and was maintained at approximately 80% humidity using a glass bubbler. Mass flow controllers (MFCs) were used to control relative humidity and the entire gas flow through the system. A LabJack U3 LV (LabJack, CO) controller was used in conjunction with a 6-way solenoid valve to facilitate an automated measurement rotation across each of the 6 jars separately over 10-minute intervals. While individual jars were in the PTR-TOF-MS measurement phase for 10 min, the remaining five jars returned to continuous flow and headspace venting.

### PTR-TOF-MS

Proton-transfer-reaction time-of-flight mass spectrometers use soft-ionization (protonation) via hydronium ions to render a single positively charged ion from volatile compounds. For example, acetic acid has a molecular monoisotopic mass/charge ratio (m/z) of 60.021, but will be detected at m/z 61.028 due to the added proton from the hydronium ion. The following protonation reaction will take place if the proton affinity of the volatile compound X is higher than that of water (Yuan et al., 2017).

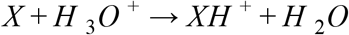

Despite being a soft-ionization method, PTR-TOF-MS can still induce fragmentation and such fragments can sometimes be used as identifiers for the mVCs of interest. For example, m/z 43.018 can be considered as an acetyl fragment originating from volatiles containing an acetyl functional group.

A PTR-TOF-MS 8000 (Ionicon Analytik, Innsbruck, Austria) was operated at a drift-tube pressure of >2.3 mbar, a drift-tube temperature of 75 oC, a field density ratio (E/N) of 120 Td, and an inlet temperature of 70 oC.

### Data Analysis

All time-series data of raw PTR-TOF-MS ion counts were normalized using the following equation

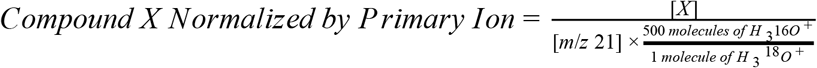

to account for the primary ion abundance. A time-series gap-filling protocol was followed using ‘dplyr’ and ‘zoo’ packages (Wickham et al., 2021; Grothendieck and Zeilleis, 2005) in R x64 3.6.3 (R Core Team, 2006) to account for 50-minute data gaps between 10-minute measurement intervals for each jar. Background signals were subtracted from sample data using empty jar measurements. Time-series data from the five soil containing jars were averaged and a spline-based time-series smoothing was performed alongside a 5-minute interval aggregation in R to reduce the effect of noise on the information partitioning (full script included in Supplementary Data. 1).

Principal components analysis (PCA) was performed on the processed time-series dataset to explore mVCs that differentiated the post wet-up phase from the preceding dry phase. R packages ‘factoextra’, ‘ggplot2’, ‘ggnewscale’ were used for PCA calculation and processing (Kassambara and Mundt, 2020; Wickham 2016; Campitelli, 2021). Each point represented in the PCA represents an mVC profile at a given 5-minute interval and colors (red for pre-wet-up, or *Dry-Phase*; and purple for post-wet-up, or *Wet-Phase*) together with gradients representing time are used to show the temporal dynamics of mVCs. Vectors representing loading scores were plotted to represent specific mVCs that contributed most to separation across the PCA axes, in addition to others that were of particular biological interest based on prior literature.

Temporal Information Partitioning Network (TIPNet)-based information partitioning was performed to go beyond synchronous metabolic connections suggested by PCA (Goodwell and Kumar, 2017). The calculations were performed using TIPNet’s graphical user interface in MATLAB (MATLAB, 2018). Mutual information (the statistical dependence between variables) over timelags from 5 to 100 minutes was calculated for every m/z pair to account for natural timelags over a series of metabolic transformations. Lagged mutual information (LMI) was further partitioned into unique (*U*), synergistic (*S*), and redundant (*R*) information to establish directionality as well as inferred causality in the relationships between variables, termed source-target relationships. This was assessed by simultaneously considering the relationships between two to three m/zs. U shows the strength of a causal relationship between two nodes (mVCs). S is a measure of new information that is obtained by using two source nodes to explain a single target node (Fig. 2). For example, trimethylamine and methanol are two different substrates that can be transformed into formaldehyde by methylotrophs and a strong synergistic connection from methanol and trimethylamine to formaldehyde could be expected. Redundant information, R was not used for this study, but is a measure of overlap in information that can be obtained by two different source nodes against a target node.

**Figure 2.**
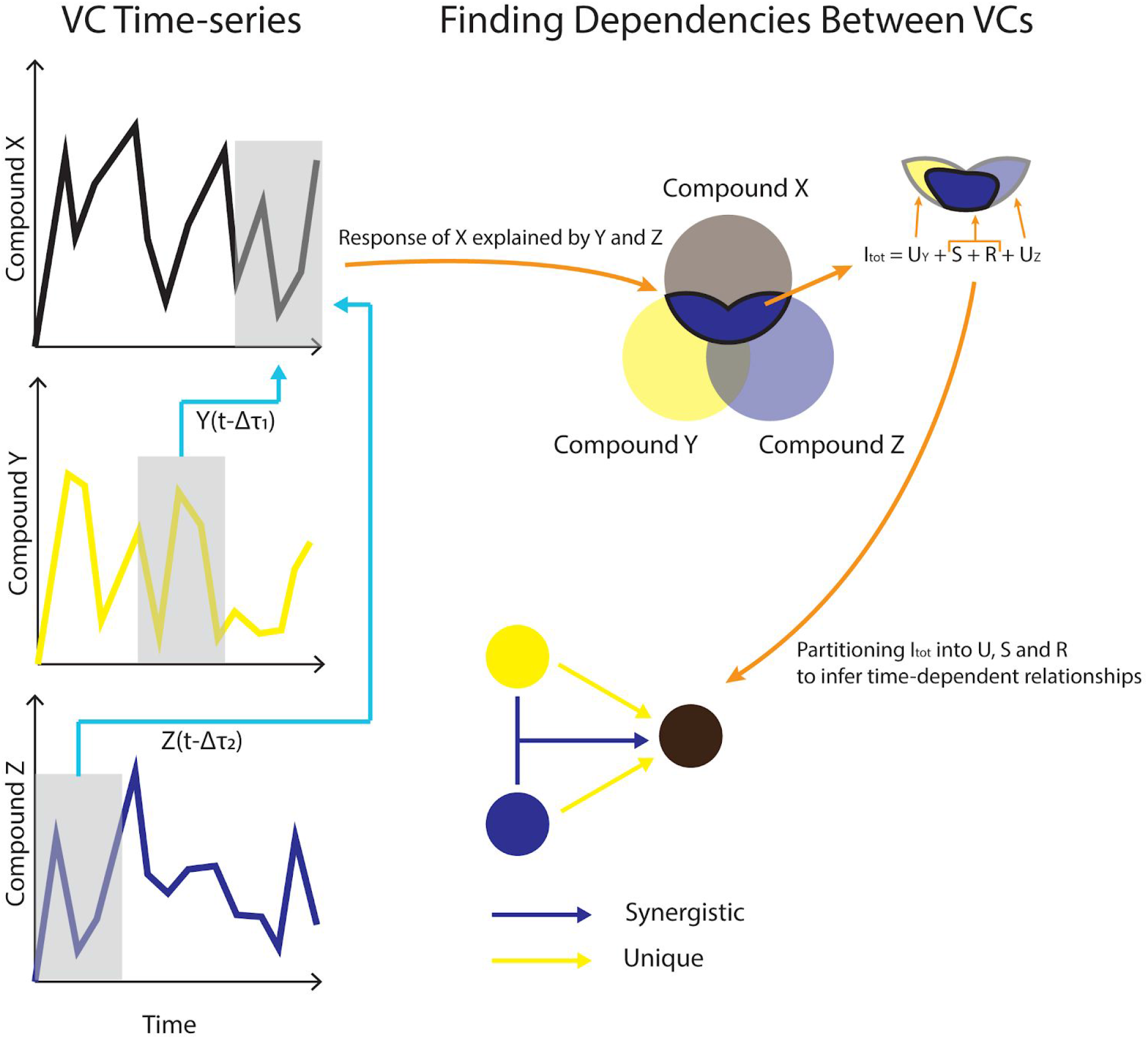
A conceptual model for using VC (Volatile Compound) time-series data to infer causality. Dependencies across different timelags in (Δτ) are used to calculate lagged mutual information. Lagged mutual information is further partitioned into unique (U), synergistic (S) and redundant (R) contributions between VCs.

Unique and synergistic information that showed statistical significance using shuffled surrogates (n=2500) were obtained from TIPNet calculations of dry and wet phases. Shuffled surrogates test for statistical significance by randomly shuffling time-series data of source time-series n number of times (Goodwell and Kumar 2017). For a simple calculation of mutual information between source node *X*(*t-*1) and target node *Y*, *X*(*t-*1) data will undergo random shuffling for ‘n’ number of times. Each U connection would imply a causal relationship between two mVCs. Each S connection would imply a metabolic relationship between the three mVCs in question (Fig. 2). For example, methanol, formaldehyde and formate are metabolites involved in methylotrophy and can form an S triad. mVCs with observed *U* and *S* relationships should be concordant in some way with metabolite relationships in known metabolic networks.

To explore this we used the KEGG reaction network and projected the top 20% (based on partitioned mutual information) of mVC pairs or triads showing significant U connections. For S connections, every source-target pair had 129 candidate mVCs to qualify as the second source contributing to the amount of explained information. Only the top mVC with the highest S information for a given source-target pair was considered as the second pair to form an S connection that later was projected onto the KEGG reaction network. KEGG pathways that included at least 2 of the inferred metabolic connections were extracted using ‘MetaboSignal’ and ‘gdata’ packages (Rodriguez-Martinez et al., 2018; Warnes et al., 2017) in R and the ‘igraph’ package (Csardi, 2005) in R was used to label the shortest paths corresponding to the *U* and *S* connections in the KEGG reaction network. The final network was created using Cytoscape 3.72 (Shannon et al., 2003). Sub-networks within the KEGG reaction network were replotted and annotated for clarity and annotated using Adobe Illustrator 2020 (24.3, Adobe Inc., available at: https://adobe.com/products/illustrator).

## 3 Results

### Principal Component Analysis of mVC Time-series

mVC profiles of the soil microcosms were initially characterized prior to and following wet-up. Time-series measurements of mVCs in the range of m/z 22-150 were obtained by PTR-TOF-MS and are included in Supplementary Data. 2. Principal Component Analysis (PCA) was performed on the entire time-series data including both dry and wet phases to identify m/z’s that characterize the wet-up response (Fig. 3). The dry phase metabolism exhibited some dynamics, oscillating around the origin of the PCA biplot. Analysis of PCA loading scores (represented as vectors overlaid on the PCA plot) indicated that the wet-up event resulted in a significant departure of metabolism and contributed most to the variance in mVC flux. In the first few hours following wet-up the mVC profile diverged abruptly from the origin before relaxing back towards the origin after the 8 day experiment. The initial divergence of metabolism upon wet-up could be ascribed to a few highly dynamic metabolites, including dimethylamine, formaldehyde, formate, followed by methanol are suggestive of methylotrophy, while the appearance of ethanol, acetaldehyde, benzaldehyde, 2,3-butanediol and diacetyl suggest the onset of fermentation and the subsequent transformation of microbial fermentation products (Jansen et al., 1984; Speckman and Collins 1968). Subsequent time points following wet-up likely experienced re-drying of the soil matrix and showed the equilibration of mVC profile back towards the origin, suggesting a disturbance-stabilization dynamic.

**Figure 3.**
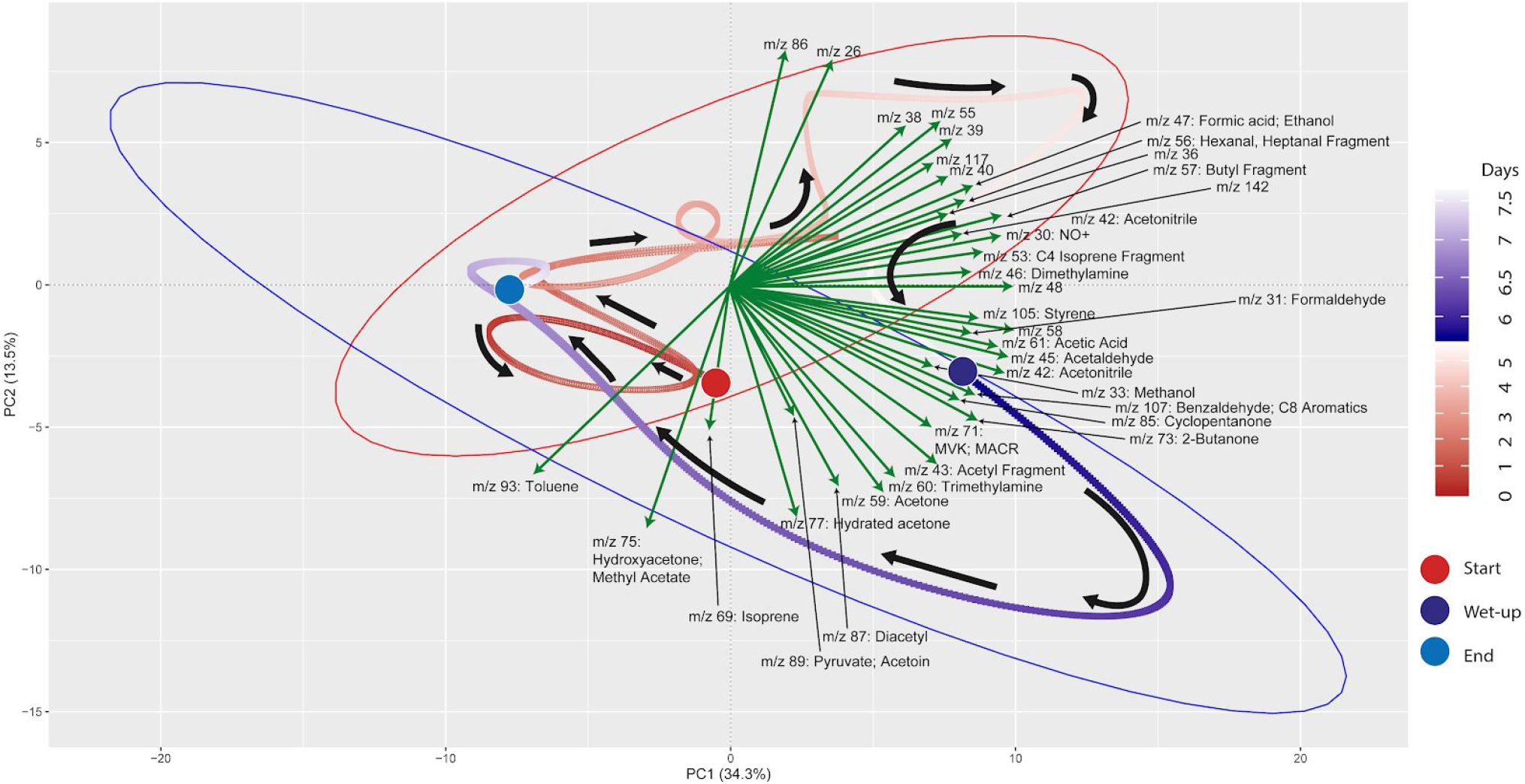
PCA (Principal Component Analysis) plot for VC (Volatile Compound) time-series data shows the effect of wet-up treatment on soil VC dynamics. Each point is VC emission profile across 5-minute time window and green arrows show specific VCs with specific m/z’s contributing to the shifts in VC profile with time and wetness. VC profile experienced the largest change during early wet-up as the time points show rapid migration towards bottom right quadrant. Subsequently, VC profile gradually stabilizes over time and returns closer to the beginning of dry measurement. Large circles mark the start and end of the incubation or start of the wet-up treatment. Pre-treatment time points are labeled in a gradient of red and post-treatment time points are labeled in a gradient of blue. Red and blue ellipses mark 95% confidence intervals of dry and wet VC profiles respectively.

### Inferring Metabolic Connections

PCA analysis provided useful information on the overall response of soil mVCs to wet-up. In order to explore the reaction dynamics responsible for observed changes in mVC emissions we performed causality inference based on Temporal Information Partitioning Networks (TIPNet). This approach allowed us to identify metabolic connections between pairs or triads of mVCs (with consideration of timelags) indicative of directional source-target relationships. In this way, each lagged mutual information between two or three mVCs was considered as a proxy for a metabolic connection and was partitioned into unique (*U*) and synergistic (*S*) information to infer causal relationships between mVCs. Unique and synergistic connections were projected onto the KEGG reaction network and yielded six sub-networks (Fig. 4) that we explore briefly below. Full sub-network information is included as a table (Supplementary Data. 3).

**Figure 4.**
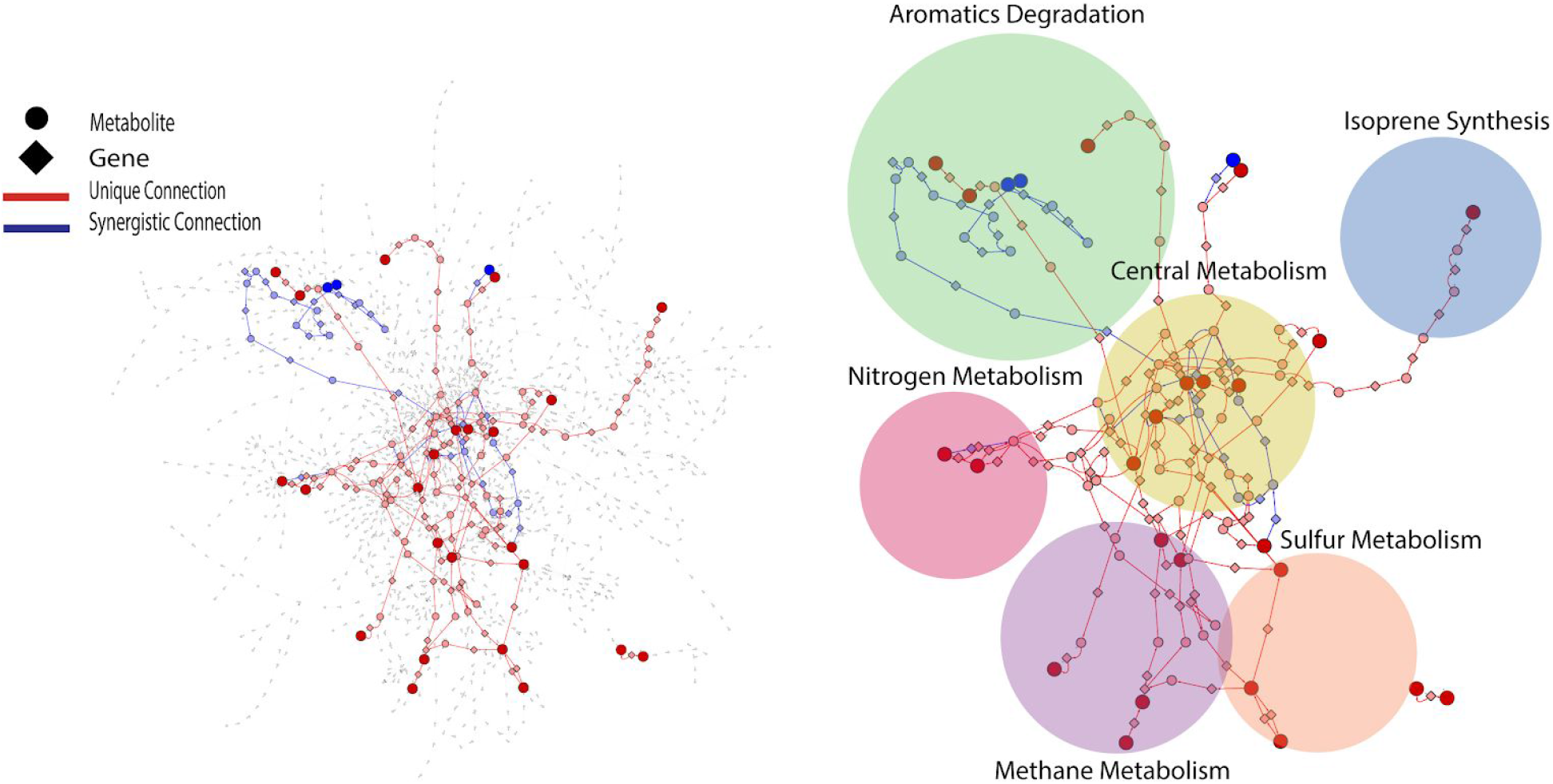
KEGG reaction network projected with metabolic connections that were tentatively detected by VC (Volatile Compound) measurement. For a given pairwise connection observed, a corresponding shortest path was mapped onto the reaction network. Each edge is a metabolite-metabolite or a metabolite-gene connection. Dark red and blue nodes are metabolites detected as VCs and light red and blue nodes are metabolites or genes that the projected shortest paths go through. The projected portion of KEGG reaction network was further divided into six sub-networks shown as circles.

### Methane and Nitrogen Metabolism

The sub-network containing KEGG pathways related to methane and nitrogen metabolism was enriched in several methylotrophic processes that pointed towards the degradation of plant litter and the possible consumption of microbial osmo/cryoprotectants upon soil re-wetting. Trimethylamine (TMA), for example is a quaternary amine known to be produced during the anaerobic degradation of glycine betaine and choline (Wang and Lee 1994), formed a synergistic connection with formaldehyde to formate. This connection is plausible given that trimethylamine can be oxidized by methylotrophs with trimethylamine dehydrogenase (EC 1.5.8.2) to dimethylamine and formaldehyde (Fig. 5). Dimethylamine also showed two unique connections to formaldehyde and formate, suggesting the resurgence of methylotrophic activity dependent on dimethylamine dehydrogenase (EC 1.5.8.1) upon wet-up. Methanol also displayed a synergistic connection with formaldehyde to formate, likely as a result of parallel methylotrophic machinery with genes, methanol dehydrogenase (EC 1.1.2.7) and formaldehyde dehydrogenase (EC 1.2.1.46) that utilize substrates derived from plant litter (i.e. pectin) de-esterification (Dorokhov et al., 2018). A unique connection identified from methanol to 2-butanone could not be projected onto an existing KEGG pathway but was included as a reaction involving an as yet unknown enzyme as 2-butanone production in soil by methylotrophs in the presence of methanol has been previously reported (Hou et al., 1979). A connection between hydroxylamine and nitrite was also included since hydroxylamine and nitrite can be produced either from ammonia oxidizing microorganisms or via methylotrophic machinery due to similarity in substrate structure and the need to detoxify resulting co-metabolites (Versantvoort et al., 2020). With this in mind, it is worth noting that ammonia is produced during TMA catabolism, and could represent a substrate source for nitrification. Metabolic connections inferred from TIPNet could also be indicated by the individual time series. Emissions of 2-butanone, trimethylamine and formaldehyde were strongly stimulated at a similar time point following wet-up, whereas methanol showed a gradual decline in emission suggestive of consumption into the aforementioned intermediates of methylotrophy.

**Figure 5.**
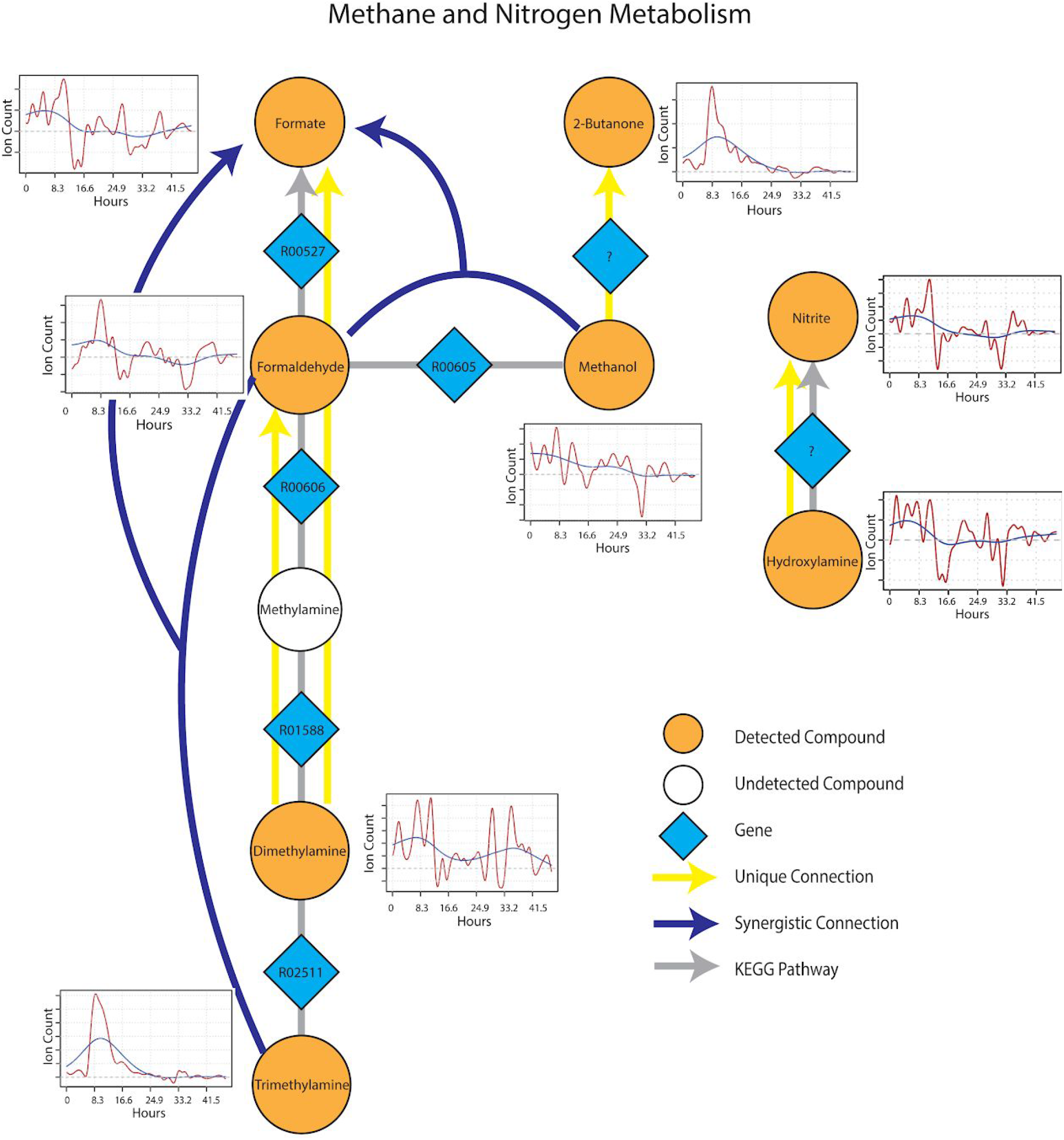
Reaction pathways that constitute methane and nitrogen metabolism sub-networks from Figure 3. Unique connections are shown in straight yellow edges and synergistic connections are shown in forked arrows. Each detected compound is labeled in orange and juxtaposed with respective time-series following wet-up. Genes responsible for each reaction are labeled in blue diamonds with respective KEGG reaction IDs. Grey edges represent KEGG pathways.

### Sulfur Metabolism

Metabolic connections in sulfur-containing mVCs were projected onto the KEGG reaction network and formed a distinct cluster containing pathways related to methylotrophy and sulfur cycling. Formaldehyde once again emerged as an important component of the sub-network being identified as a target mVC with sources including dimethyl sulfide (DMS) and methanethiol (Fig. 6), projectable on KEGG pathways driven by dimethyl sulfide monooxygenase (EC 1.14.13.131) and methanethiol oxidase (EC 1.8.3.4) respectively. In particular, methanethiol also formed a synergistic connection to formaldehyde with hydrogen sulfide as the second source, which consolidates the role of methanethiol oxidase (EC 1.8.3.4) in the observed connection. Two non-methylotrophic pathways were also shown by methanethiol and hydrogen sulfide. Methanethiol formed a unique connection to dimethyl sulfide and this connection projected onto a *mddA* methyltransferase pathway in KEGG (EC 2.1.1.334). Hydrogen sulfide was uniquely connected to acetate and the connection was projected onto a *CysO* cysteine synthase pathway with acetate as its target mVC (EC 2.5.1.47). Time-series data of both methanethiol and DMS differed from those of pure carbon mVCs as they showed a more gradual increase and peaked at a later time point with patterns showing an apparent co-variance. This co-variance that was exclusive to sulfur-containing mVCs suggested not only the metabolic connectivity of the two compounds but also the metabolic progression of the soil microbial community that shows evidence of mostly oxidative and some fermentative carbon and nitrogen metabolism followed by a reduced sulfur-based one, possibly suggesting a change in redox conditions.

**Figure 6.**
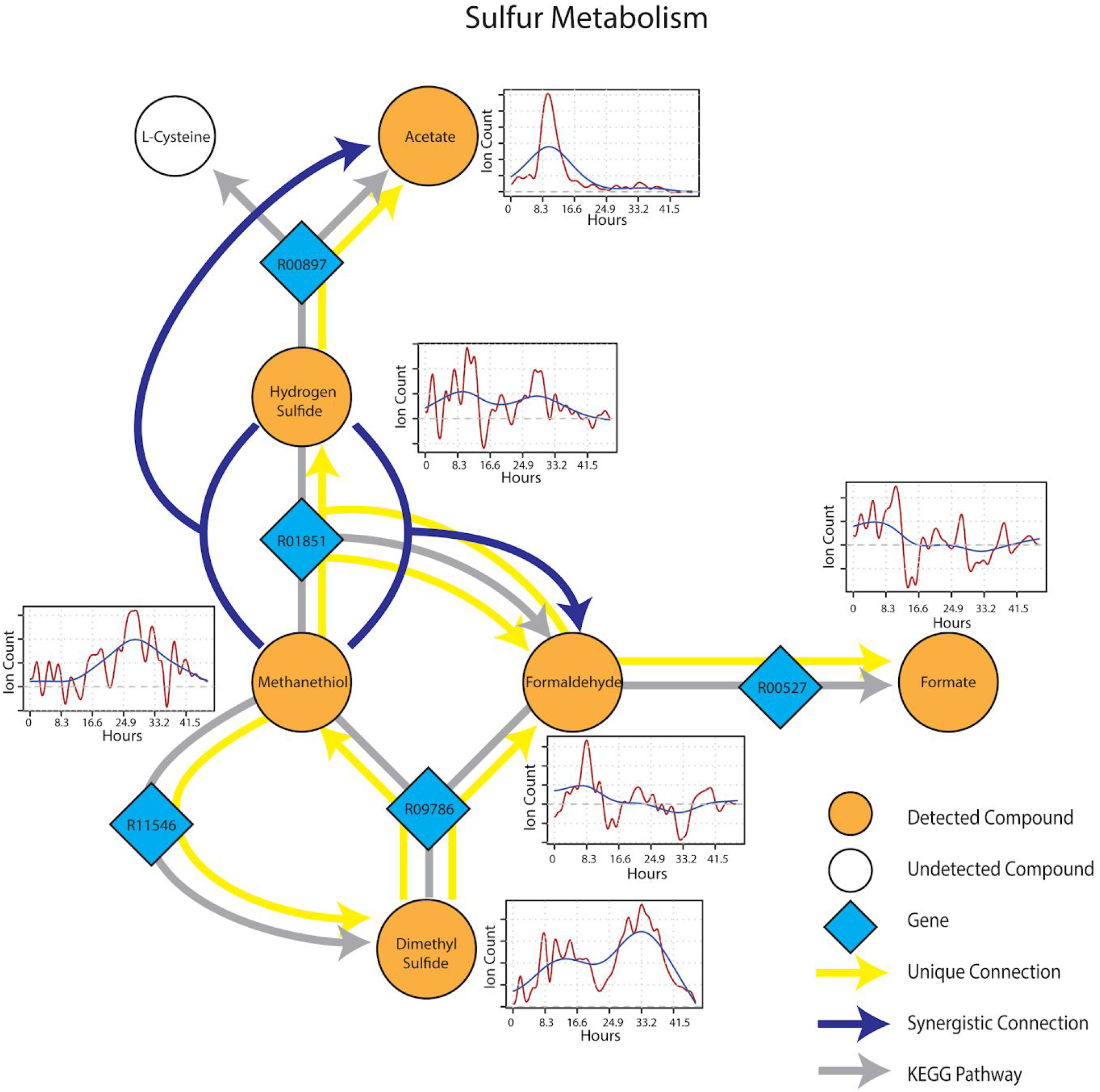
Reaction pathways that constitute sulfur metabolism sub-network from Figure 3. Unique connections are shown in straight yellow edges and synergistic connections are shown in forked arrows. Each detected compound is labeled in orange and juxtaposed with respective time-series following wet-up. Genes responsible for each reaction are labeled in blue diamonds with respective KEGG reaction IDs. Grey edges represent KEGG pathways.

### Central Metabolism and Isoprene

After wet-up, processes related to fermentation and lower redox potential were predominant in the central metabolism and isoprene sub-networks. Pyruvate showed a unique connection to diacetyl and two different synergistic connections to acetate and 2,3-butanediol paired with methanol (Fig. 7). These connections were projected onto fermentation pathways from KEGG involving acetolactate decarboxylase (EC 4.1.1.5), acetate kinase (EC 2.7.2.1) and 2,3-butanediol dehydrogenase (EC 1.1.1.4) respectively. Ethanol appeared as both a unique source and a target mVC connecting to and from acetaldehyde via pathways likely driven by alcohol dehydrogenase (EC 1.1.1.1). A unique connection from isoprene to acetate was also included (although it could not be projected onto the current KEGG reaction network) based on the hypothesis from previous studies that isoprene is oxidized through beta-oxidation pathway with an unknown enzyme (McGenity et al., 2018). Time-series trendlines of pyruvate and diacetyl showed apparent co-variance with 2,3-butanediol also showing an emission spike at a similar time point in accordance with the 2,3-butanediol fermentation from pyruvate to diacetyl and 2,3-butanediol. The time-series of pyruvate and acetaldehyde also shared a similar pattern indicative of a close connection within the KEGG metabolic network.

**Figure 7.**
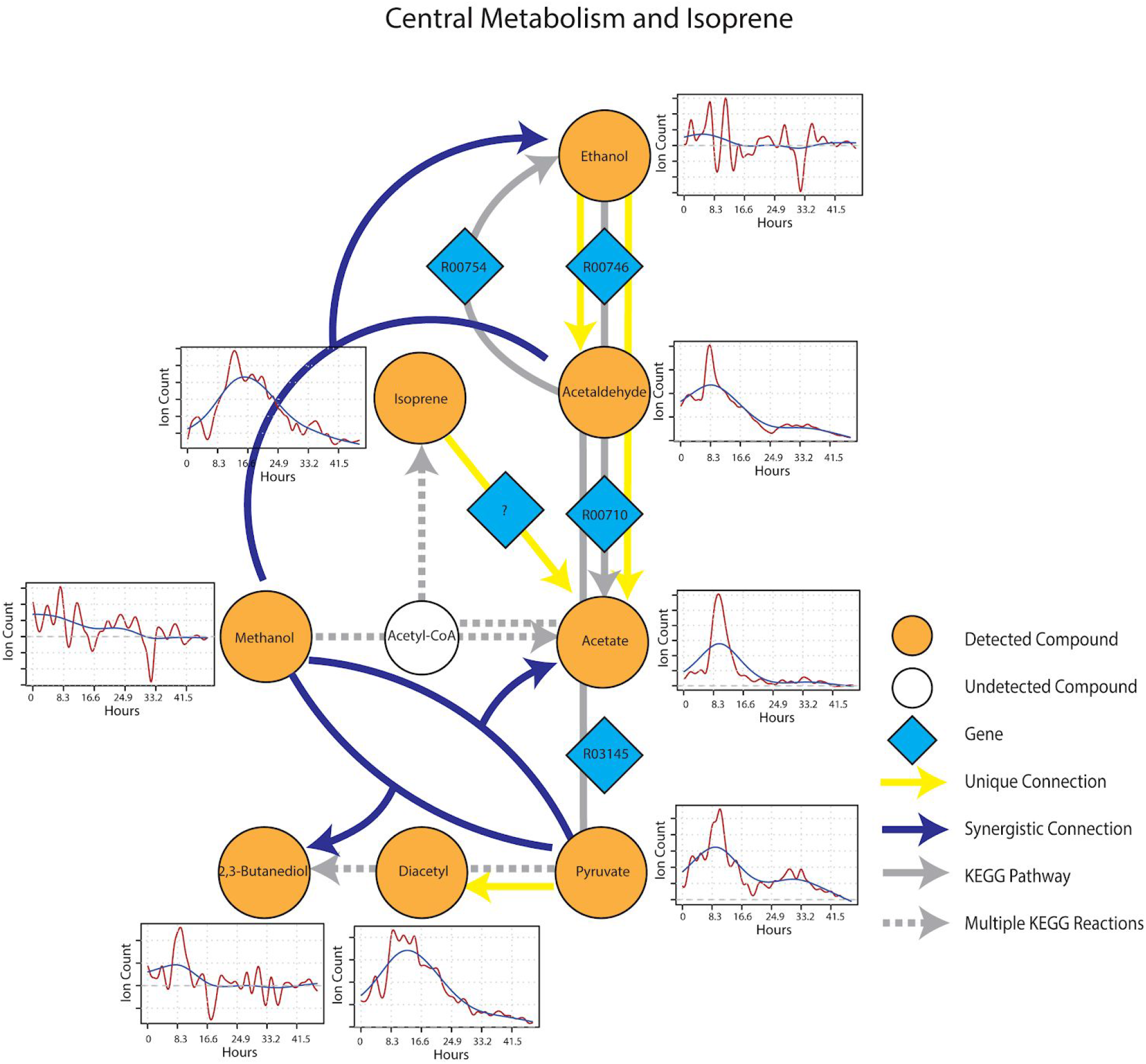
Reaction pathways that constitute central metabolism and isoprene sub-network from Figure 3. Unique connections are shown in straight yellow edges and synergistic connections are shown in forked arrows. Each detected compound is labeled in orange and juxtaposed with respective time-series following wet-up. Genes responsible for each reaction are labeled in blue diamonds with respective KEGG reaction IDs. Grey edges represent KEGG pathways.

Overall, we saw a great deal of concordance between predicted metabolic connections based on mVCs and those with prior evidence as represented in the KEGG reaction network. This provided a data-driven approach to non-destructively monitor the metabolic trajectories of soil microbial communities in response to soil re-wetting in a manner similar to a snowmelt event, and provides predictions as to which metabolic pathways and enzymes may be stimulated under these conditions.

## 4 Discussion

Microbial volatile compounds (mVCs) have been used as metabolic fingerprints in multiple disciplines including epidemiology, microbial ecology, and environmental microbiology (Palma et al., 2018; McNeal and Herbert, 2009; Misztal et al., 2018) and have demonstrated utility as tractable and non-destructive windows into microbial metabolism. Most prior studies focused on the detection of individual mVCs while treating the emission profile itself as a fingerprint. Our approach took advantage of semi-continuous time-series of mVC fluxes to establish causal linkages between metabolites using information theory analyses. This allowed us to use the observed emission profile to explain the biological state of the system, namely soil. Our goal was to leverage on mVCs as a window into the specific biological pathways constituting the biogeochemical state of soil. We explored the possibility of real-time characterization of soil microbiome metabolic processes by using time-series mVC data from PTR-TOF-MS measurements. Ordination of mVC profiles as they transitioned from the initial comparatively dry condition of soil that is often characteristic of that beneath a snowpack, to the wet-up phase akin to the onset of snowmelt, showed a rapid and distinct change in the trajectory of microbial metabolism. This wet-up driven activation of soil microbiome metabolism was ephemeral with the metabolic state (reflected as PCA coordinates) generally returning towards the origin within 4 days after wet-up (Fig. 3). The infiltration of snowmelt is known to induce a pronounced increase in soil microbial growth and metabolism (e.g. Lipson et al., 2002; Sorensen et al., 2020). As expected, we observed an accumulation during wet-up of mVCs such as diacetyl and butanol known to be produced under oxygen limiting conditions (Seewald et al., 2010). We also observed a dynamic emission of mVCs possibly related to methylotrophy (e.g. methylamines, methanol) and saw further evidence of the onset of fermentation (such as ethanol, acetaldehyde, benzaldehyde, and 2,3-butanediol).

Re-wetting of soil could result in mVC pulses by purely abiotic factors such as chemo-desorption caused by water addition (Thibaud et al., 1993). This uncertainty required an approach where the biological relevance of observed mVCs could be assessed in the context of biological pathways. Directed connections between mVCs were obtained based on lagged mutual information (LMI) and were used to construct a putative reaction network that could be projected onto a known biological network (KEGG in this case) to evaluate the biological relevance of the mVC fluxes. Information partitioning of LMI was used to calculate unique and synergistic connections that showed pair and triad relationships to further narrow down the type of microbial processes contributing to respective connections. In this regard, two synergistic connections, methanol & formaldehyde to formate and trimethylamine(TMA) & formaldehyde to formate (Fig. 5) strongly indicated that methylotrophic activity was stimulated by wet-up. Prior qNMR data from soils at a nearby site showed high abundance of methanol during snowmelt (unpublished data). The source of methanol remains uncertain but may be related to the de-esterification of plant litter (Dorokhov et al., 2018) and consequently increased the activity of methylotrophs present in soil. However, a rapid increase in trimethylamine (TMA) emission was also observed upon wet-up. TMA is a substrate for methylotrophs and it has been previously observed to originate from the anaerobic degradation quaternary amines such as choline and glycine betaine by microbes (Wang et al., 1994). Glycine betaine is shown to function both as both an osmo- and cryo-protectant in microbes and plants (Chattopadhyay et al., 2002; Xing and Rajashekar, 2001), thus it is possible that upon soil wet-up, microbial lysis or active export of osmo/cryoprotectants are transformed to TMA for eventual consumption by methylotrophs (Sun et al., 2019). Other mVCs that we observed during early wet-up such as acetone, methanol and acetaldehyde have previously been observed in soil following precipitation and are commonly regarded as volatile signatures of plant litter decomposition and support a hypothesis that litter decomposition was stimulated by water addition (Schade and Goldstein 2001; Schade and Goldstein 2002; Schink and Zeikus 2007; Niemenmaa et al. 2008; Gray et al. 2010).

This approach of identifying unique and synergistic connections also allows interactions between carbon and nutrient cycles to be postulated. For example, following the initial pulse of mVCs during wet-up, rapid production of TMA potentially from anaerobic degradation of quaternary amines was soon matched by consumption at a similar rate (Fig. 5) which could be driven by the activation of methylotrophs. The synergistic connection of TMA & formaldehyde -> formate (Fig. 5) supports that TMA was consumed by methylotrophs as formaldehyde is an important byproduct in TMA degradation via trimethylamine dehydrogenase often found in methylotrophs (Sun et al., 2019). Dimethylamine (DMA) also displayed a unique connection to formaldehyde and TMA was likely degraded into DMA leading to sequential extraction of formaldehyde from available methylamines. Full oxidation of methylamine can yield an ammonium ion that can be directly utilized by methylotrophs or released to the extracellular space becoming available for cross-feeding by nearby microbial communities (Taubert et al., 2017). Although we could not measure methylamine emission via PTR-TOF-MS due to high background from molecular oxygen (m/z 32), the rapid consumption of TMA via methylotrophic pathway suggests an equivalent release of ammonium ions as a result of methylamine oxidation. This hypothesis aligns with the results of another study from our group on nearby soils that showed a rapid increase in microbial biomass N early upon snowmelt initialization followed by a population crash, a pulse of nitrate in parallel with an increase in nitrifier abundance (Sorensen et al., 2020). Interestingly, a unique connection from hydroxylamine to nitrite was observed and could be a result of a known detoxification process managed by either methylotroph or nitrifier hydroxylamine oxidoreductase, possibly as a result of the release of ammonium ions from methylamine oxidation.

Combined, these results support a hypothesis that the change in soil water content and redox conditions similar to the conditions of snowmelt, stimulated methylotrophic activity by providing methylamines from anaerobic decomposition of quaternary amines, and consequently leading to an increase in ammonium ion pool to stimulate further autotrophic or heterotrophic microbial growth. Further studies are required to confirm these possibilities.

While early wet-up was characterized by rapid and diverse production of mVCs, later timepoints generally showed lower diversity and intensity of mVCs being emitted. Sulfur-containing mVCs, methanethiol and dimethyl sulfide (DMS), showed a gradual increase in emission as TMA emission disappeared (Fig. 6). Both methanethiol and DMS have been shown to occur in anaerobic soil (Stotzky et al., 1976). The sources remain unclear, but previous studies have confirmed that methanethiol can be produced as a result of methionine degradation, whereas DMS is produced via methylation of methanethiol and appears to be a widespread phenomenon across soils (Carrión et al., 2017). Regardless of the source of these sulfur-containing mVCs, they are also methylotrophic substrates like TMA and indeed showed unique and synergistic connections to formaldehyde (Fig. 4). It appears that the stimulation of methylotrophic activity may be partitioned into two stages; an early, but ephemeral methylotrophy driven by TMA and degraded amines, and a later stage that provides DMS and methanethiol.

As presented, we propose that information partitioning of time-series data from PTR-TOF-MS can be used to build a metabolic narrative of soil microbiome metabolism and offers a practical means of diagnosing the activation of major metabolic pathways following perturbations. This approach is amenable to other diagnostic purposes such as evaluating soil redox transitions and estimating the significance of anaerobic microsites. In this regard, diacetyl, 2,3-butanediol and other mVCs that are known fermentation byproducts are useful markers and metabolic connections, such as pyruvate to 2,3-butanediol, and acetaldehyde to ethanol can be computed to further verify the biological pathways suggested by the presence of the mVCs detected. Further, this approach can also be used to generate hypotheses regarding unexplored pathways. For example, in our study, a unique connection, methanol to 2-butanone, suggests that this metabolic transformation is occurring, and while soil methanol-utilizers have been shown to produce 2-butanone, the responsible enzymes have yet to be identified (Hou et al., 1979). Similarly, isoprene to acetate is a unique connection that supports a general, but unverified assumption that isoprene is degraded via beta-oxidation pathway to supply carbon to the central metabolism (McGenity et al., 2018; Carrión et al., 2018). Such hypotheses that can be generated by mVC analysis provide room for complementary use of other methods such as stable isotope tracing and probing.

In this proof-of-principle study, we explored an information theory approach where mVC time-series are analyzed to discover unique and synergistic metabolite-to-metabolite connections that are projectable to KEGG reaction networks. We showed that this approach can be useful in providing a metabolic narrative that coincides with the results from other studies. A number of sophisticated metabolomic approaches exist with higher fidelity, however a necessity to destructively sample makes the assessment of microbial dynamics and metabolic couplings difficult to interpret and precludes true time-series measurements. Continuous or semi-continuous time-series data also allows directional connections to be computed from lagged mutual information and its partitioned constituents such as unique and synergistic information. Overall, we propose that mVCs, especially when combined with online measurement such as PTR-MS, represent an attractive approach to “remotely sense” soil microbiome metabolism.

## Acknowledgements

This work was supported by the Laboratory Directed Research and Development Program of Lawrence Berkeley National Laboratory under U.S. Department of Energy Contract No. DE-AC02-05CH11231. JK was also supported by the UC Berkeley Honors Program, Sponsored Projects for Undergraduate Research (SPUR). We thank the members of the LBNL Watershed Function Scientific Focus Area (SFA), particularly Dr. Rosemary Carrol, Dr. Jeffrey Deems for obtaining soil samples and Dr. Luca Peruzzo for providing data on soil hydraulic properties.

## Supplementary Materials

All data and code are included as supplementary materials.

Supplementary Data 1: All R scripts and softwares used for data processing

Supplementary Data 2: Time-series measurements of mVCs in the range of m/z 22-150 obtained by PTR-TOF-MS.

Supplementary Data 3: Table showing full sub-network relationships from information entropy analysis of mVC dynamics.

## References

Barnard, R. L., Blazewicz, S. J., Firestone, M. K. (2020). Rewetting of soil: Revisiting the origin of soil CO2 emissions. Soil Biology and Biochemistry. 147, 107819. doi: 10.1016/j.soilbio.2020.107819.

Birch, H. F. (1958). The Effect of Soil Drying On Humus Decomposition and Nitrogen Availability. Plant and Soil. 10, 9–31.

Blazewicz, S., Hungate, B., Koch, B., Nuccio, E., Morrissey, E., Brodie, E., et al. (2020). Taxon-specific microbial growth and mortality patterns reveal distinct temporal population responses to rewetting in a California grassland soil. The ISME Journal. 14 doi: 10.1038/s41396-020-0617-3.

Brooks, P., Williams, M., Schmidt, S. (1998). Inorganic Nitrogen and Microbial Biomass Dynamics Before and During Spring Snowmelt. Biogeochemistry. 43, 1–15. doi: 10.1023/A:1005947511910.

Campitelli, E. (2021). ggnewscale: Multiple Fill and Colour Scales in ‘ggplot2’. R package version 0.4.5. https://CRAN.R-project.org/package=ggnewscale.

Carrión, O., Larke-Mejía, N. L., Gibson, L., Farhan Ul Haque, M., Ramiro-García, J., McGenity, T. J., et al. (2018). Gene probing reveals the widespread distribution, diversity and abundance of isoprene-degrading bacteria in the environment. Microbiome. 6, 219. doi: 10.1186/s40168-018-0607-0.

Carrión, O., Pratscher, J., Curson, A. R. J., Williams, B. T., Rostant, W. G., Murrell, J. C., et al. (2017). Methanethiol-dependent dimethylsulfide production in soil environments. The ISME Journal. 11, 2379–90. doi: 10.1038/ismej.2017.105.

Chattopadhyay, M. K. (2002). The cryoprotective effects of glycine betaine on bacteria. Trends in Microbiology. 10, 311. doi: 10.1016/S0966-842X(02)02395-8.

Csardi, G., Nepusz, T. (2005). The Igraph Software Package for Complex Network Research. InterJournal. Complex Systems, 1695.

Goodwell, A. E., Jiang, P., Ruddell, B. L., Kumar, P. (2020). Debates—Does Information Theory Provide a New Paradigm for Earth Science? Causality, Interaction, and Feedback. Water Resources Research. 56, e2019WR024940. doi: https://doi.org/10.1029/2019WR024940.

Goodwell, A. E., Kumar, P. (2017). Temporal Information Partitioning Networks (TIPNets): A process network approach to infer ecohydrologic shifts. Water Resources Research. 53, 5899–919. doi: https://doi.org/10.1002/2016WR020218.

Gray, C. M., Monson, R. K., Fierer, N. (2010). Emissions of volatile organic compounds during the decomposition of plant litter. Journal of Geophysical Research: Biogeosciences. 115 doi: https://doi.org/10.1029/2010JG001291.

Grothendieck, G., Zeileis, A. (2005). zoo: S3 Infrastructure for Regular and Irregular Time Series. Journal of Statistical Software. 14 doi: 10.18637/jss.v014.i06.

Hou, C. T., Patel, R., Laskin, A. I., Barnabe, N., Marczak, I. (1979). Microbial oxidation of gaseous hydrocarbons: production of methyl ketones from their corresponding secondary alcohols by methane- and methanol-grown microbes. Appl Environ Microbiol. 38, 135–42. doi: 10.1128/AEM.38.1.135-142.1979.

Jansen, N. B., Flickinger, M. C., Tsao, G. T. (1984). Production of 2,3-butanediol from D-xylose by Klebsiella oxytoca ATCC 8724. Biotechnology and Bioengineering. 26, 362–9. doi: https://doi.org/10.1002/bit.260260411.

Jiang, P., Kumar, P. (2019). Information transfer from causal history in complex system dynamics. Phys Rev E. 99, 012306. doi: 10.1103/PhysRevE.99.012306.

Kanehisa, M., Goto, S. (2000). KEGG: Kyoto Encyclopedia of Genes and Genomes. Nucleic Acids Res. 28, 27–30. doi: 10.1093/nar/28.1.27.

Kassambara, A., and Mundt, F. (2020). factoextra: Extract and Visualize the Results of Multivariate Data Analyses. R package version 1.0.7. https://CRAN.R-project.org/package=factoextra.

Lipson, D. A., Schadt, C. W., Schmidt, S. K. (2002). Changes in soil microbial community structure and function in an alpine dry meadow following spring snow melt. Microb Ecol. 43, 307–14. doi: 10.1007/s00248-001-1057-x.

MATLAB. (2018). 9.7.0.1190202 (R2019b). Natick, Massachusetts: The MathWorks Inc.

McGenity, T., Crombie, A. T., Murrell, J. (2018). Microbial cycling of isoprene, the most abundantly produced biological volatile organic compound on Earth. The ISME Journal. doi: 10.1038/s41396-018-0072-6.

McNeal, K. S., Herbert, B. E. (2009). Volatile Organic Metabolites as Indicators of Soil Microbial Activity and Community Composition Shifts. Soil Science Society of America Journal. 73, 579–88. doi: 10.2136/sssaj2007.0245.

Misztal, P. K., Lymperopoulou, D. S., Adams, R. I., Scott, R. A., Lindow, S. E., Bruns, T., et al. (2018). Emission Factors of Microbial Volatile Organic Compounds from Environmental Bacteria and Fungi. Environ Sci Technol. 52, 8272–82. doi: 10.1021/acs.est.8b00806.

Netzker, T., Shepherdson, E., Zambri, M., Elliot, M. (2020). Bacterial Volatile Compounds: Functions in Communication, Cooperation, and Competition. Annual Review of Microbiology. 74 doi: 10.1146/annurev-micro-011320-015542.

Niemenmaa, O., Uusi-Rauva, A., Hatakka, A. (2008). Demethoxylation of [O14CH3]-labelled lignin model compounds by the brown-rot fungi Gloeophyllum trabeum and Poria (Postia) placenta. Biodegradation. 19, 555–65. doi: 10.1007/s10532-007-9161-3.

Palma, Susana I. C. J., Traguedo, A. P., Porteira, A., Frias, M. J., Gamboa, H., Roque, A. (2018). Machine learning for the meta-analyses of microbial pathogens’ volatile signatures. Scientific Reports. doi: 10.1038/s41598-018-21544-1.

Peters, A., Iden, S. C., and Durner, W. (2015). Revisiting the simplified evaporation method: Identification of hydraulic functions considering vapor, film and corner flow. Journal of Hydrology, 527:531–542.

Placella, S. A., Brodie, E. L., Firestone, M. K. (2012). Rainfall-induced carbon dioxide pulses result from sequential resuscitation of phylogenetically clustered microbial groups. PNAS. 109, 10931–6.

Placella, S., Firestone, M. (2013). Transcriptional Response of Nitrifying Communities to Wetting of Dry Soil. Applied and environmental microbiology. 79 doi: 10.1128/AEM.00404-13.

R Core Team. (2006). A language and environment for statistical computing. Computing. 1 doi: 10.1890/0012-9658(2002)083[3097:CFHIWS]2.0.CO;2.

Rodriguez-Martinez, A., Ayala, R., Posma, J. M., Dumas, M. (2018). Exploring the Genetic Landscape of Metabolic Phenotypes with MetaboSignal. Current Protocols in Bioinformatics. 61, 14.14.1,14.14.13. doi: https://doi.org/10.1002/cpbi.41.

Schade, G. W., Goldstein, A. H. (2001). Fluxes of oxygenated volatile organic compounds from a ponderosa pine plantation. Journal of Geophysical Research: Atmospheres. 106, 3111–23. doi: https://doi.org/10.1029/2000JD900592.

Schade, G. W., Goldstein, A. H. (2002). Plant physiological influences on the fluxes of oxygenated volatile organic compounds from ponderosa pine trees. Journal of Geophysical Research: Atmospheres. 107, ACH 2–8. doi: https://doi.org/10.1029/2001JD000532.

Schindler, U., Durner, W., von Unold, G., Mueller, L., and Wieland, R. (2010a). The evaporation method: Extending the measurement range of soil hydraulic properties using the air-entry pressure of the ceramic cup. Journal of Plant Nutrition and Soil Science, 173(4):563–572.

Schindler, U., Durner, W., von Unold, G., and Müller, L. (2010b). Evaporation Method for Measuring Unsaturated Hydraulic Properties of Soils: Extending the Measurement Range. Soil Science Society of America Journal, 74(4):1071–1083.

Schink, B., Zeikus, J. (2007). Microbial methanol formation: A major end product of pectin metabolism. Current Microbiology. doi: 10.1007/BF02605383.

Seewald, M., Singer, W., Knapp, B., Franke-Whittle, I., Hansel, A., Insam, H. (2010). Substrate-induced volatile organic compound emissions from compost-amended soils. Biology and Fertility of Soils. doi: 10.1007/s00374-010-0445-0.

Shannon, P. (2003). Cytoscape: A Software Environment for Integrated Models of Biomolecular Interaction Networks. Genome research. 13, 2498–504. doi: 10.1101/gr.1239303.

Sims, G. K., Ellsworth, T. R., Mulvaney, R. L. (1995). Microscale determination of inorganic nitrogen in water and soil extracts. Communications in Soil Science and Plant Analysis. 26, 303–16. doi: 10.1080/00103629509369298.

Sorensen, P. O., Bhatnagar, J. M., Christenson, L., Duran, J., Fahey, T., Fisk, M. C., et al. (2019). Roots Mediate the Effects of Snowpack Decline on Soil Bacteria, Fungi, and Nitrogen Cycling in a Northern Hardwood Forest. Front Microbiol. 10 doi: 10.3389/fmicb.2019.00926.

Sorensen, P. O., Beller, H. R., Bill, M., Bouskill, N. J., Hubbard, S. S., Karaoz, U., et al. (2020). The Snowmelt Niche Differentiates Three Microbial Life Strategies That Influence Soil Nitrogen Availability During and After Winter. Front Microbiol. 11 doi: 10.3389/fmicb.2020.00871.

Speckman, R. A., Collins, E. B. (1968). Diacetyl biosynthesis in Streptococcus diacetilactis and Leuconostoc citrovorum. J Bacteriol. 95, 174–80. doi: 10.1128/JB.95.1.174-180.1968.

Steltzer, H., Landry, C., Painter, T. H., Anderson, J., Ayres, E. (2009). Biological consequences of earlier snowmelt from desert dust deposition in alpine landscapes. Proc Natl Acad Sci U S A. 106, 11629–34. doi: 10.1073/pnas.0900758106.

Stotzky, G., Schenck, S., Papavizas, G. C. (1976). Volatile Organic Compounds and Microorganisms. CRC Critical Reviews in Microbiology. 4, 333–82. doi: 10.3109/104084176091023

Sun, J., Mausz, M. A., Chen, Y., Giovannoni, S. J. (2019). Microbial trimethylamine metabolism in marine environments. Environmental Microbiology. 21, 513–20. doi: https://doi.org/10.1111/1462-2920.14461.

Taubert, M., Grob, C., Howat, A. M., Burns, O. J., Pratscher, J., Jehmlich, N., et al. (2017). Methylamine as a nitrogen source for microorganisms from a coastal marine environment. Environmental Microbiology. 19, 2246–57. doi: https://doi.org/10.1111/1462-2920.13709.

Thibaud, C., Erkey, C., Akgerman, A. (1993). Investigation of the effect of moisture on the sorption and desorption of chlorobenzene and toluene from soil. Environ Sci Technol. 27, 2373–80. doi: 10.1021/es00048a010.

Tyc, O., Song, C., Dickschat, J. S., Vos, M., Garbeva, P. (2017). The Ecological Role of Volatile and Soluble Secondary Metabolites Produced by Soil Bacteria. Trends in Microbiology. 25, 280–92. doi: 10.1016/j.tim.2016.12.002.

van Genuchten, M. T. (1980). A Closed-form Equation for Predicting the Hydraulic Conductivity of Unsaturated Soils. Soil Science Society of America Journal, 44(5):892.

van Genuchten, M. T. and Nielsen, D. R. (1985). On Describing and Predicting the Hydraulic Properties of Unsaturated Soil. Annales Geophysicae, 3:615–628.

Versantvoort, W., Pol, A., Jetten, M. S. M., van Niftrik, L., Reimann, J., Kartal, B., et al. (2020). Multiheme hydroxylamine oxidoreductases produce NO during ammonia oxidation in methanotrophs. PNAS. 117, 24459–63.

Wang, X., Lee, C. (1994). Sources and distribution of aliphatic amines in salt marsh sediment. Organic Geochemistry. 22, 1005–21. doi: 10.1016/0146-6380(94)90034-5.

Warnes, G., Bolker, B., Gorjanc, G., Grothendieck, G., Korosec, A., Lumley, T., MacQueen, D., Magnusson, A., Rogers, J. (2017). gdata: Various R Programming Tools for Data Manipulation. R package version 2.18.0. https://CRAN.R-project.org/package=gdata.

Wickham, H. (2016). ggplot2: Elegant Graphics for Data Analysis. Springer. ISBN 978-3-319-24277-4, https://ggplot2.tidyverse.org.

Wickham, H., François, R., Henry, L., and Müller, K. (2021). dplyr: A Grammar of Data Manipulation. R package version 1.0.4. https://CRAN.R-project.org/package=dplyr.

Wipf, S. (2010). Phenology, Growth, and Fecundity of Eight Subarctic Tundra Species in Response to Snowmelt Manipulations. Plant Ecology. 207, 53–66.

Xing, W., Rajashekar, C. B. (2001). Glycine betaine involvement in freezing tolerance and water stress in Arabidopsis thaliana. Environmental and Experimental Botany. 46, 21–8. doi: 10.1016/S0098-8472(01)00078-8.

