## Supplementary Data 1 for "Measurement of Volatile Compounds for Real-time Analysis of Soil Microbial Metabolic Response to Simulated Snowmelt": Readme.txt

Supplementary Data. 1

Contents

Inlet1.txt: Raw mVC time-series data from first soil replicate 
Inlet2.txt: Raw mVC time-series data from second soil replicate 
Inlet3.txt: Raw mVC time-series data from third soil replicate 
Inlet4.txt: Raw mVC time-series data from fourth soil replicate 
Inlet5.txt: Raw mVC time-series data from fifth soil replicate 
Inlet6.txt: Raw mVC time-series data from blank jar
Source_Code.txt: Entire source code used for data processing in R
