## Supplementary Data 2 for "Measurement of Volatile Compounds for Real-time Analysis of Soil Microbial Metabolic Response to Simulated Snowmelt"

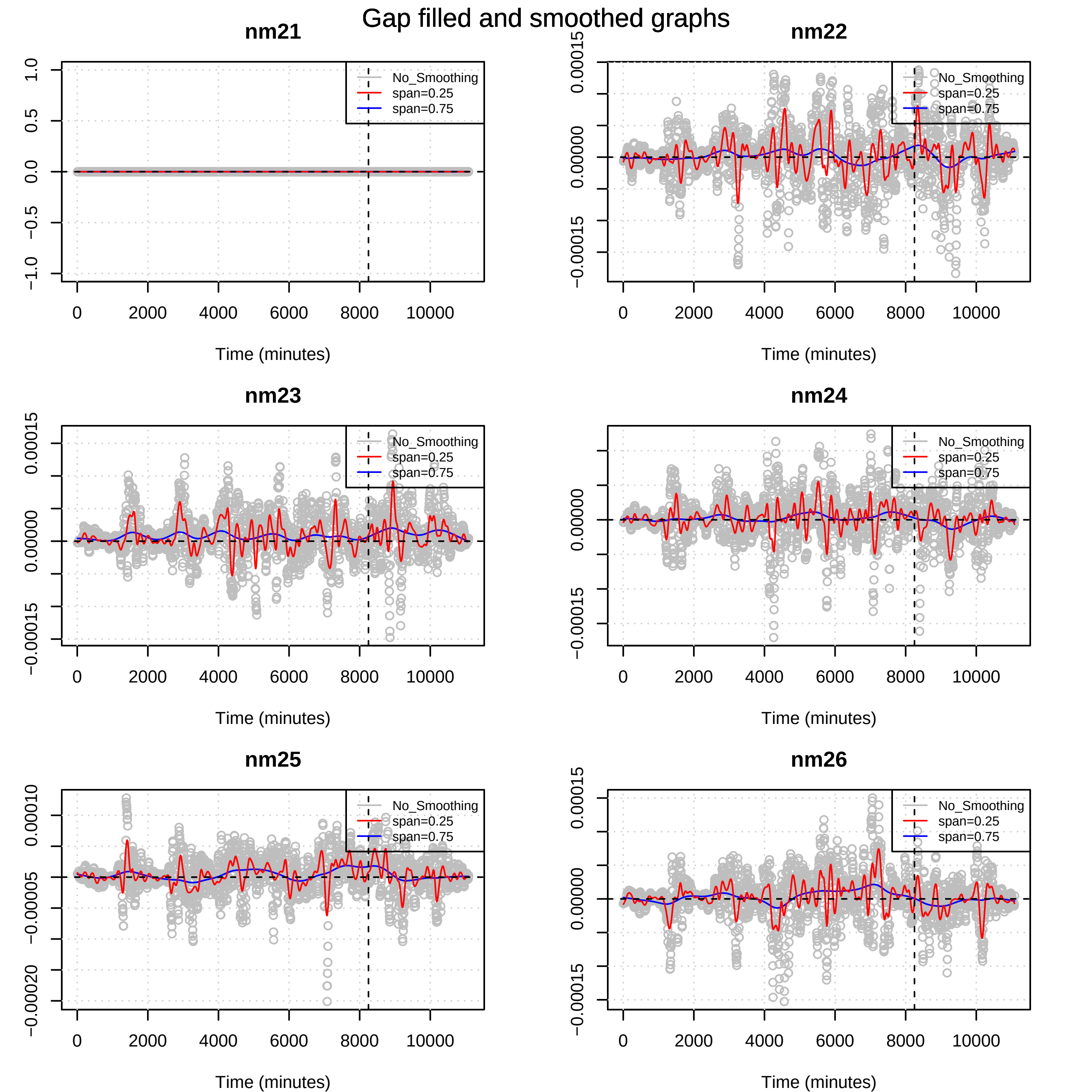

Gap filled and smoothed graphs
nm21
nm22
nm23
nm24
nm25
nm26
No_Smoothing
span=0.25
span=0.75
Time (minutes)
0
2000
4000
6000
8000
10000
1.0
0.5
0.0
−0.5
−1.0
0.00015
0.00000
−0.00015
0.00010
−0.00005
−0.00020

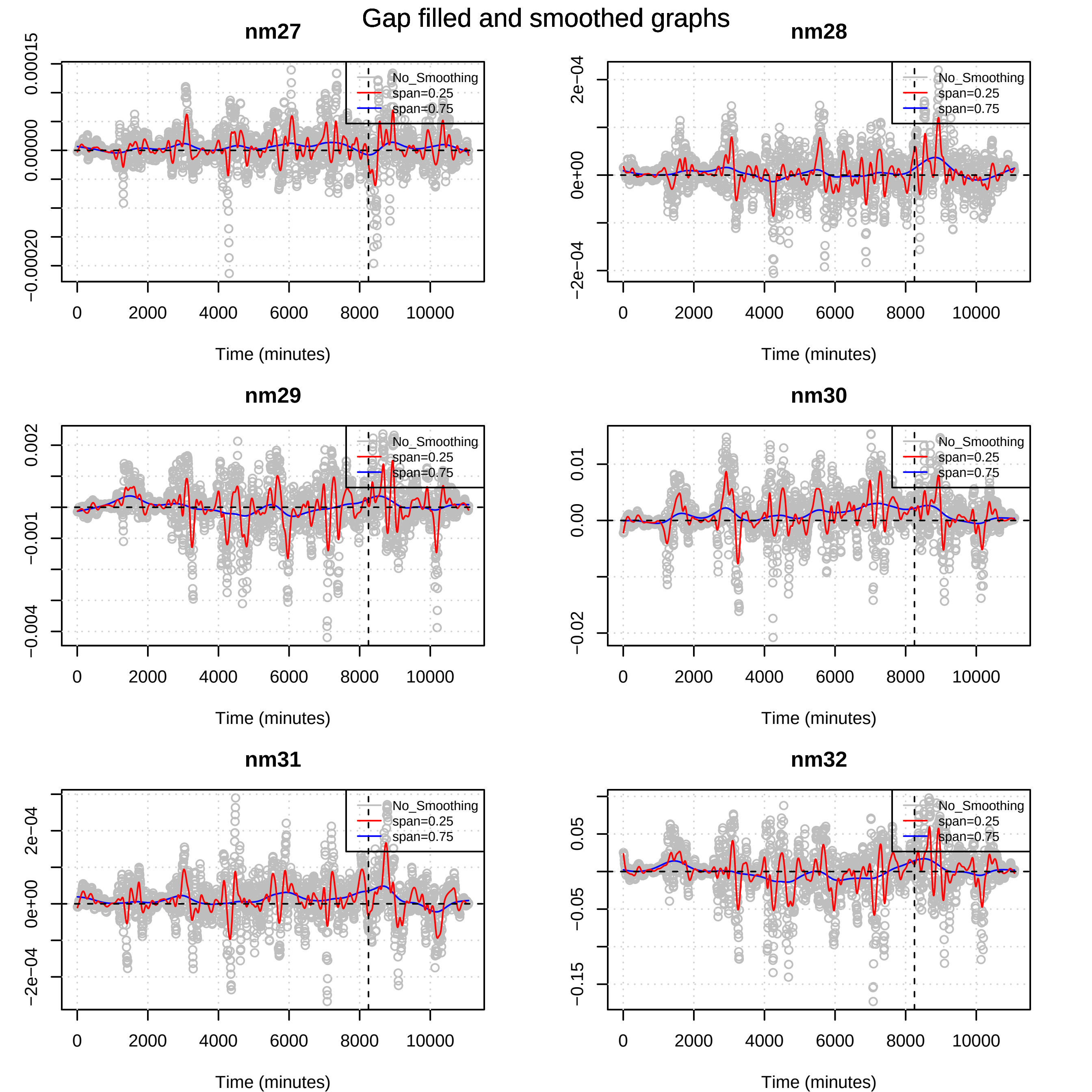

Gap filled and smoothed graphs
nm27
nm28
nm29
nm30
nm31
nm32
No_Smoothing
span=0.25
span=0.75
Time (minutes)
0
2000
4000
6000
8000
10000
0.00015
0.00000
−0.00020
2e−04
0e+00
−2e−04
0.002
−0.001
−0.004
0.01
0.00
−0.02
0.05
−0.05
−0.15

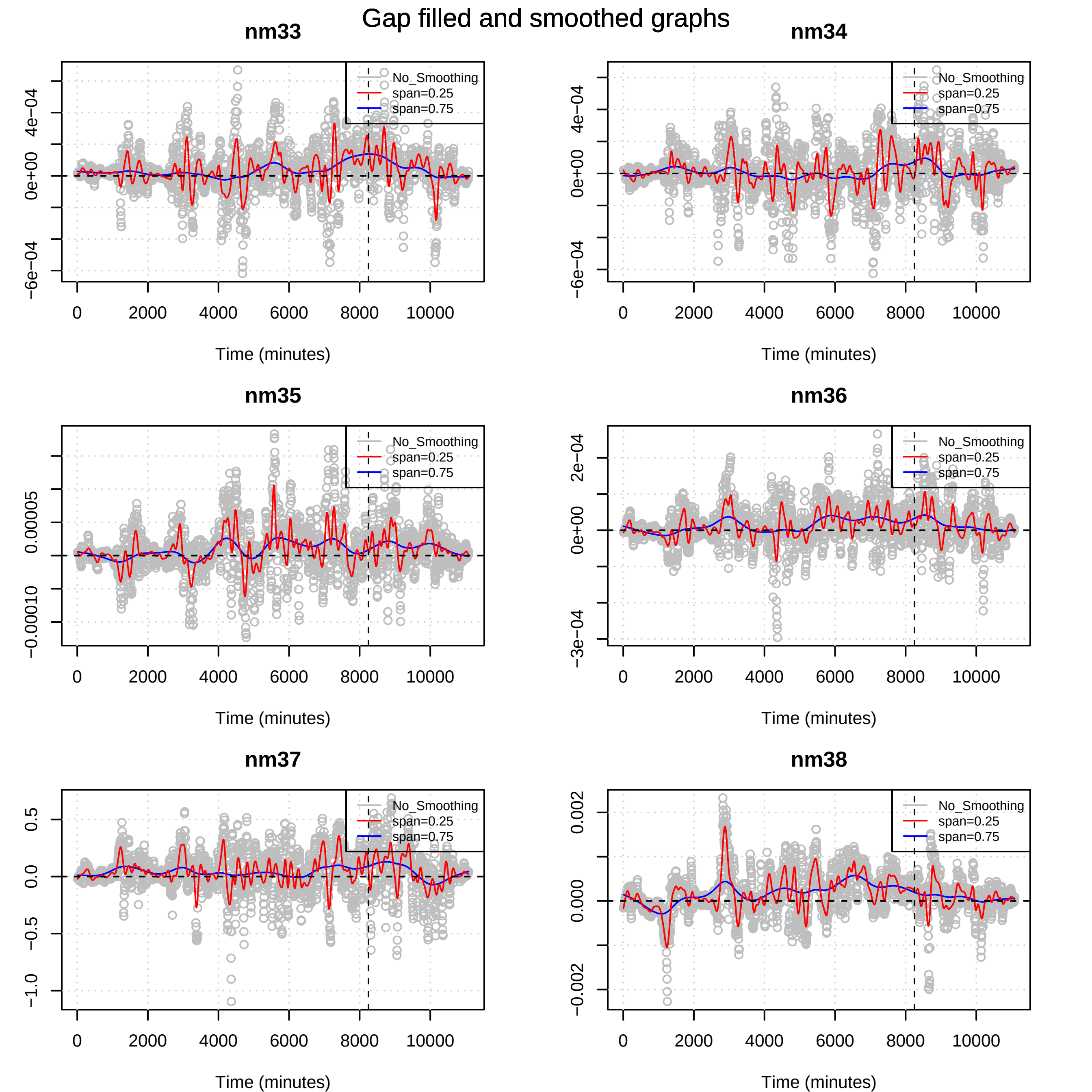
Gap filled and smoothed graphs
nm33
nm34
nm35
nm36
nm37
nm38
No_Smoothing
span=0.25
span=0.75
Time (minutes)
0
2000
4000
6000
8000
10000
−6e−04
0e+00
4e−04
−0.00010
0.00005
−3e−04
2e−04
−1.0
−0.5
0.0
0.5
−0.002
0.000
0.002

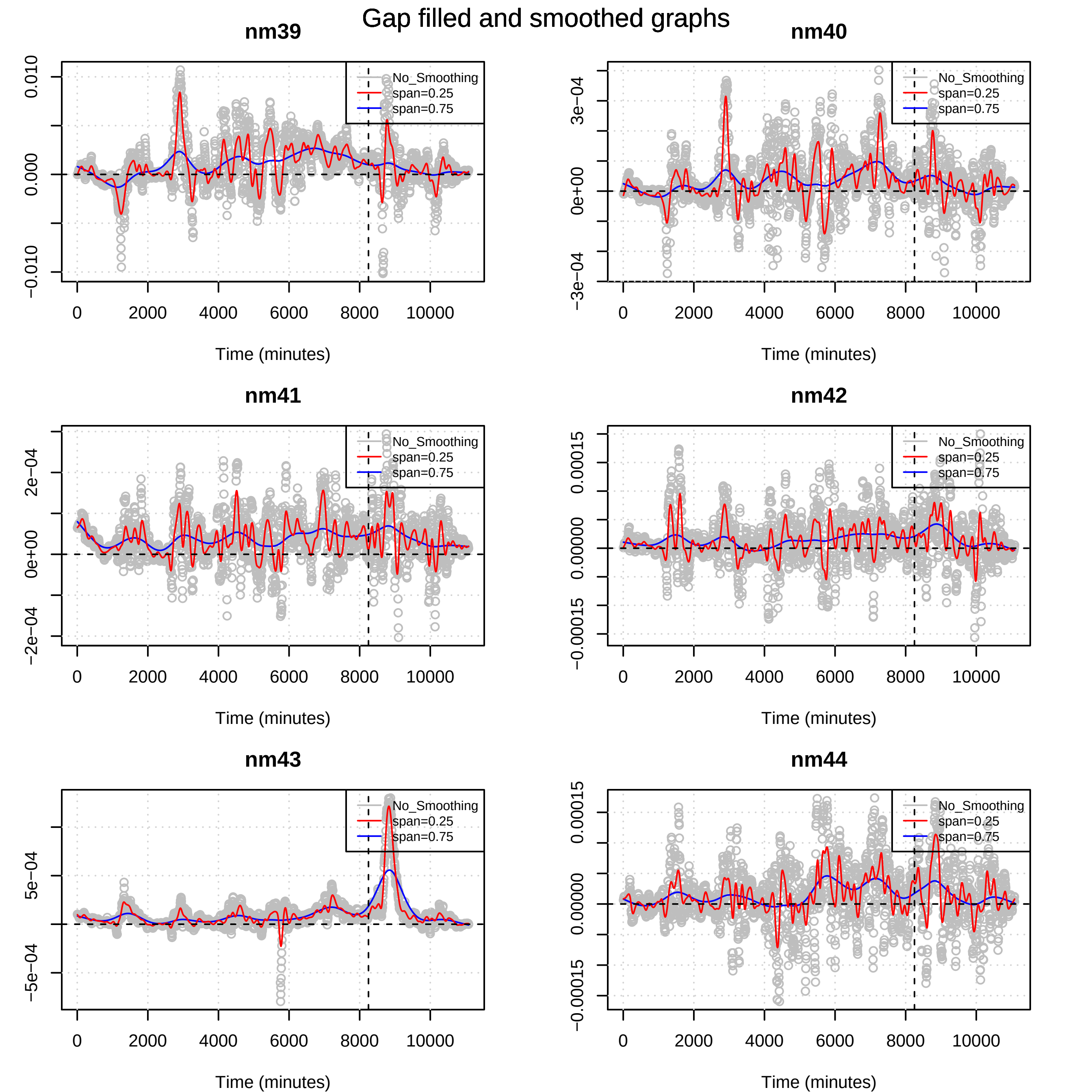

Gap filled and smoothed graphs
nm39
nm40
nm41
nm42
nm43
nm44
No_Smoothing
span=0.25
span=0.75
Time (minutes)
0
2000
4000
6000
8000
10000
0.010
0.000
−0.010
3e−04
0e+00
−3e−04
2e−04
−2e−04
0.00015
0.00000
−0.00015
5e−04
−5e−04

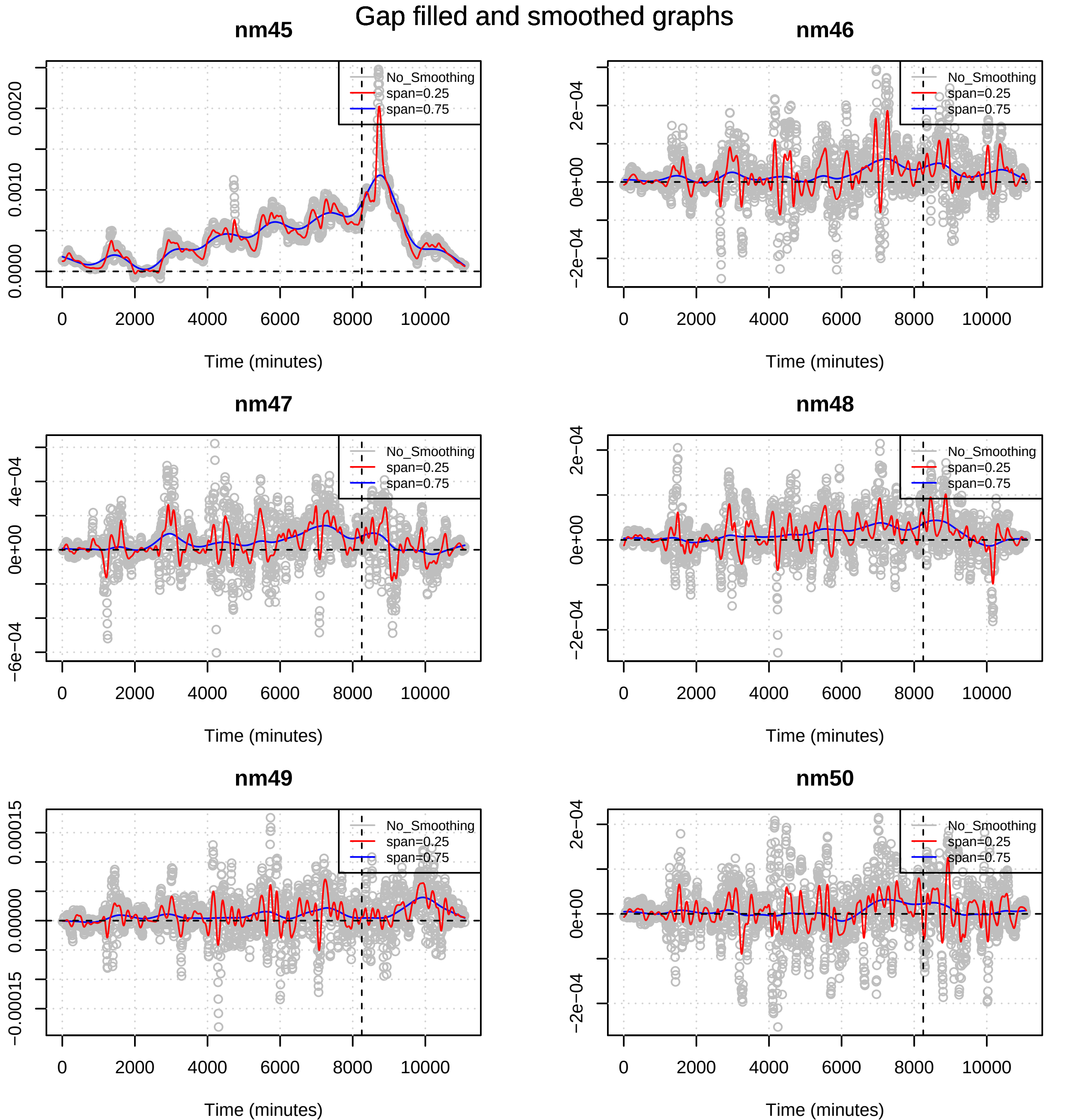

Gap filled and smoothed graphs
nm45
nm46
nm47
nm48
nm49
nm50
No_Smoothing
span=0.25
span=0.75
0.0000
0.0010
0.0020
2e−04
0e+00
−2e−04
4e−04
−6e−04
0.00015
0.00000
−0.00015
0
2000
4000
6000
8000
10000
Time (minutes)

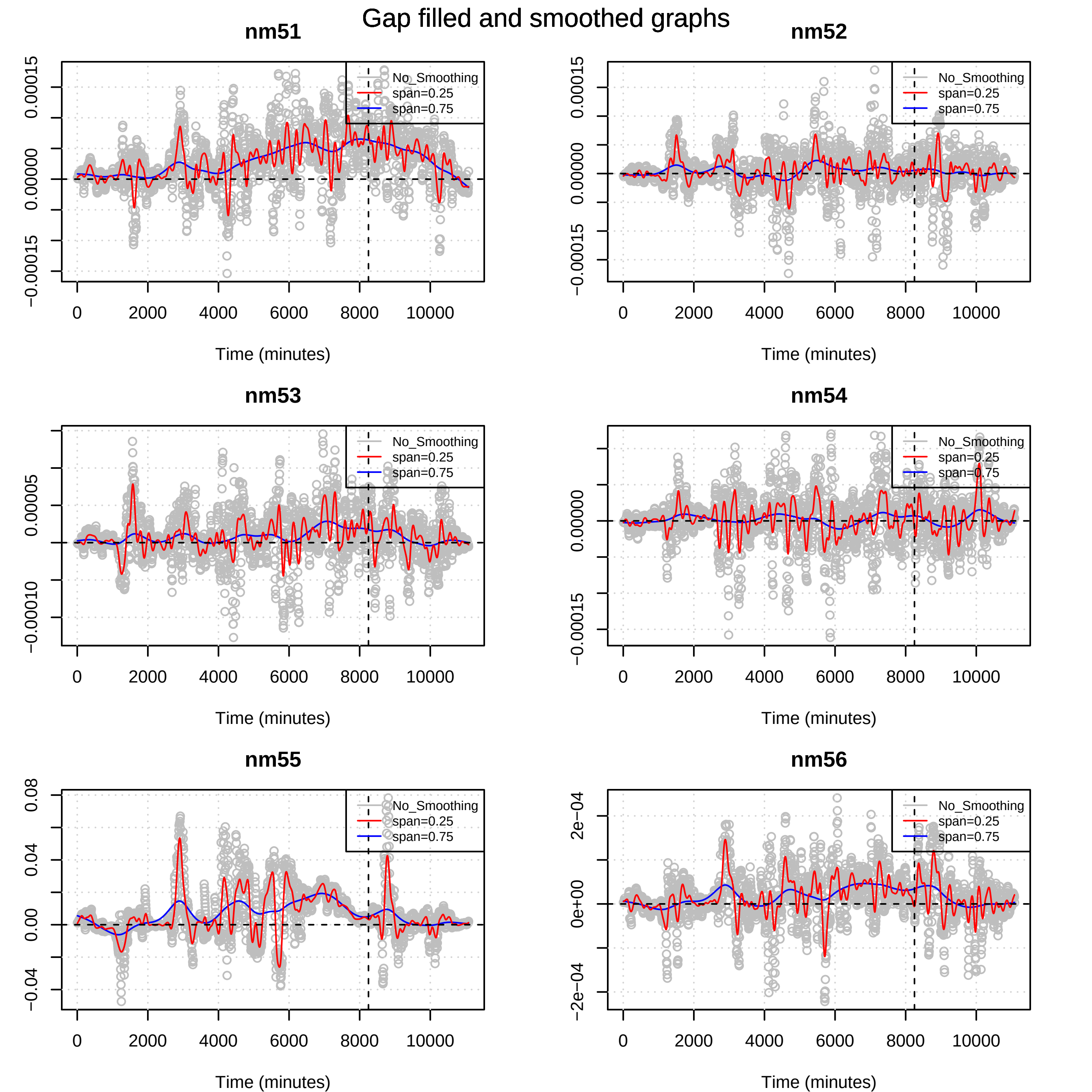
Gap filled and smoothed graphs
nm51
nm52
nm53
nm54
nm55
nm56
No_Smoothing
span=0.25
span=0.75
Time (minutes)
0
2000
4000
6000
8000
10000
0.00015
0.00000
−0.00015
0.00005
−0.00010
0.08
0.04
0.00
−0.04
2e−04
0e+00
−2e−04

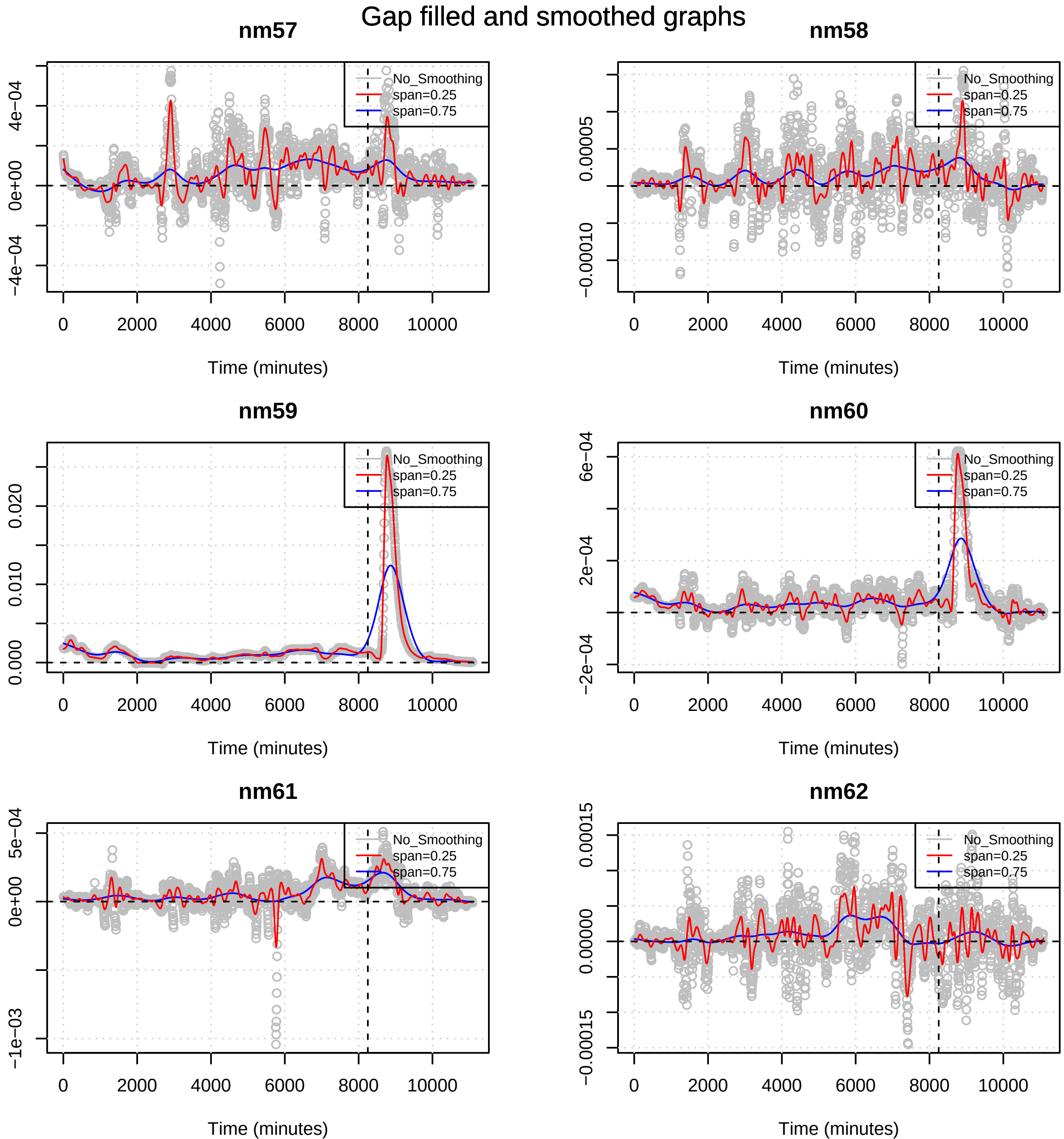

Gap filled and smoothed graphs
nm57
nm58
nm59
nm60
nm61
nm62
No_Smoothing
span=0.25
span=0.75
Time (minutes)
0
2000
4000
6000
8000
10000
4e−04
0e+00
−4e−04
0.00005
−0.00010
0.000
0.010
0.020
6e−04
2e−04
−2e−04
5e−04
0e+00
−1e−03
0.00015
0.00000
−0.00015

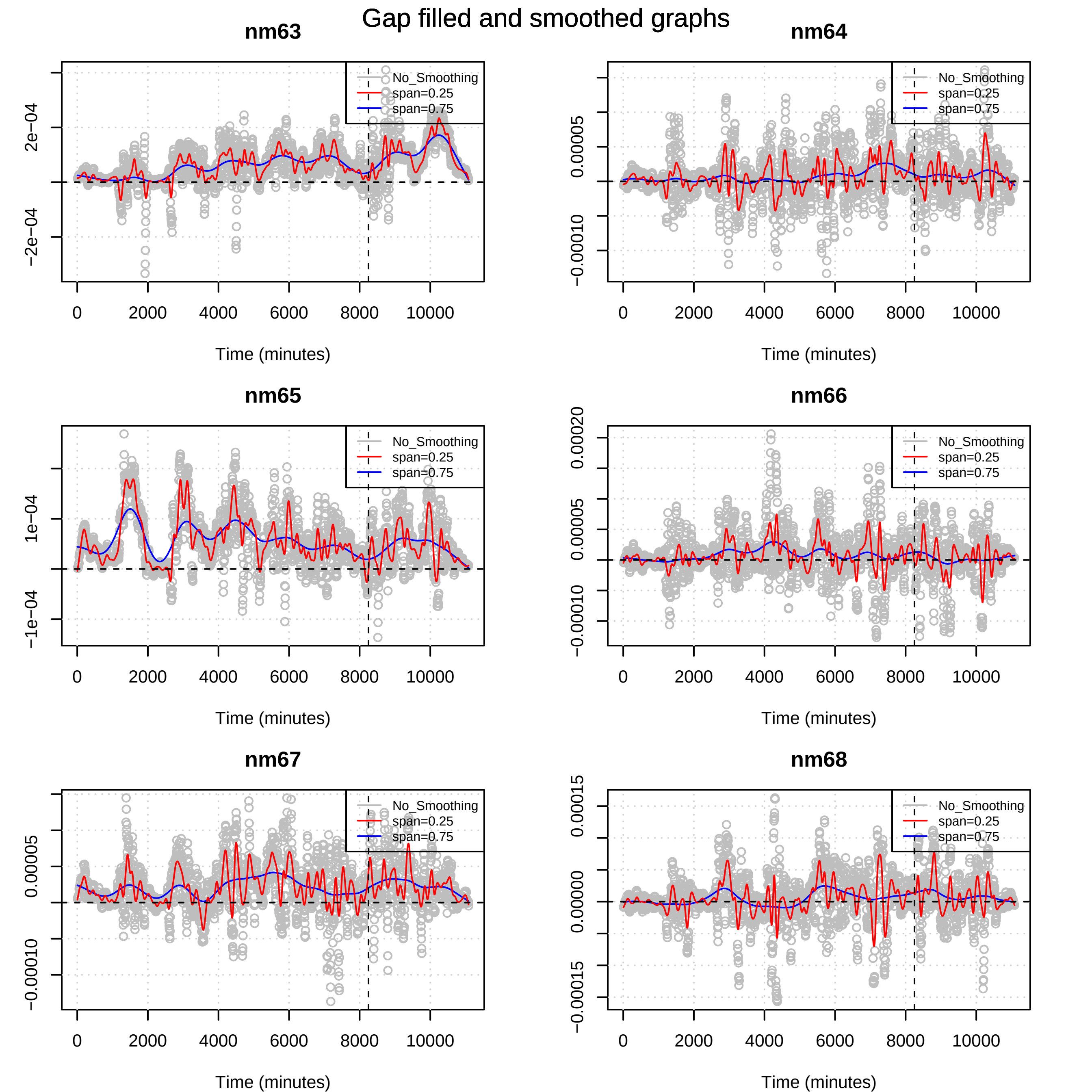

Gap filled and smoothed graphs
nm63
nm64
nm65
nm66
nm67
nm68
No_Smoothing
span=0.25
span=0.75
Time (minutes)
0
2000
4000
6000
8000
10000
2e−04
−2e−04
1e−04
−1e−04
0.00005
−0.00010
0.00020
0.00015
0.00000
−0.00015

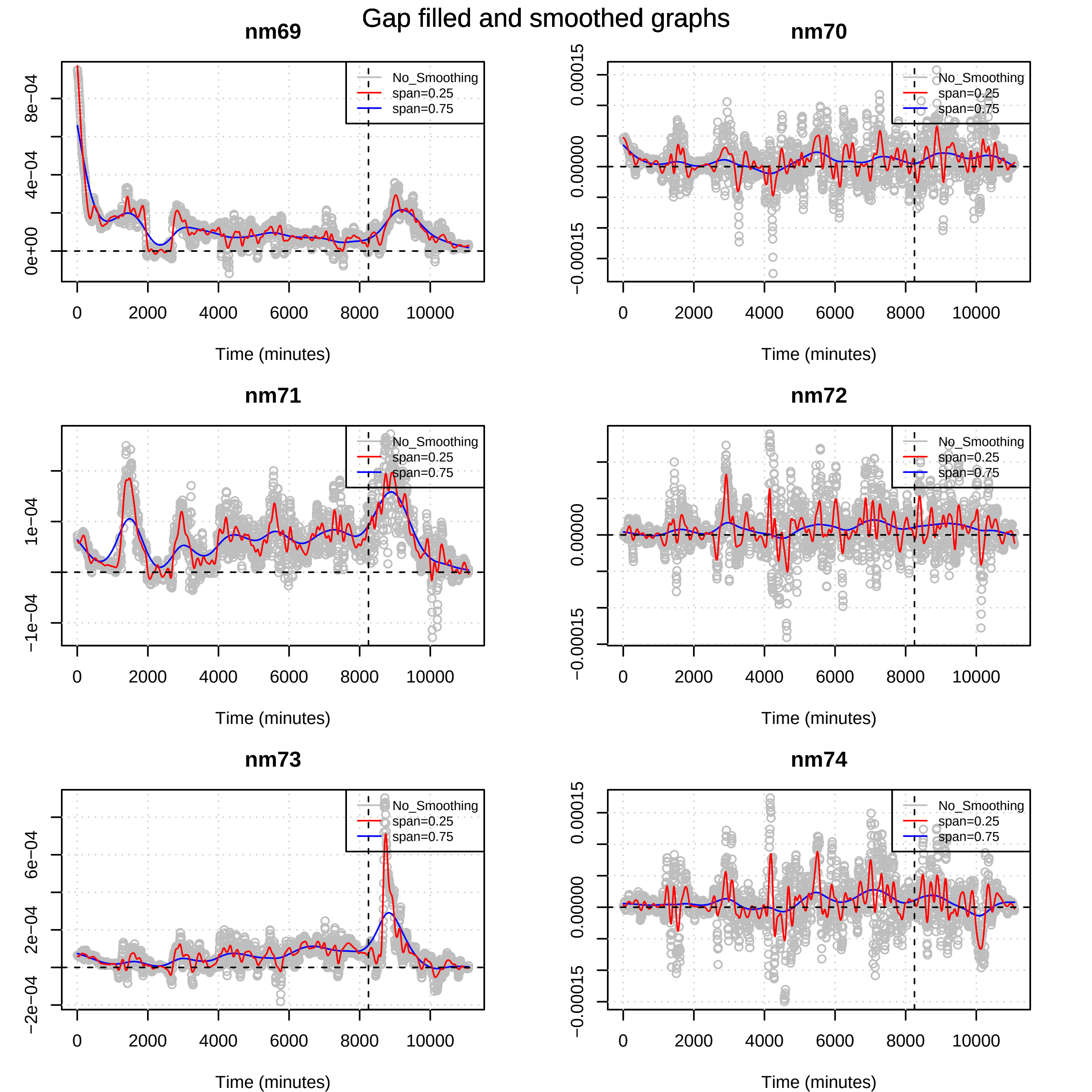
Gap filled and smoothed graphs
nm69
nm70
nm71
nm72
nm73
nm74
No_Smoothing
span=0.25
span=0.75
Time (minutes)
0
2000
4000
6000
8000
10000
0e+00
4e−04
8e−04
−0.00015
0.00000
0.00015
−1e−04
1e−04
−2e−04
2e−04
6e−04

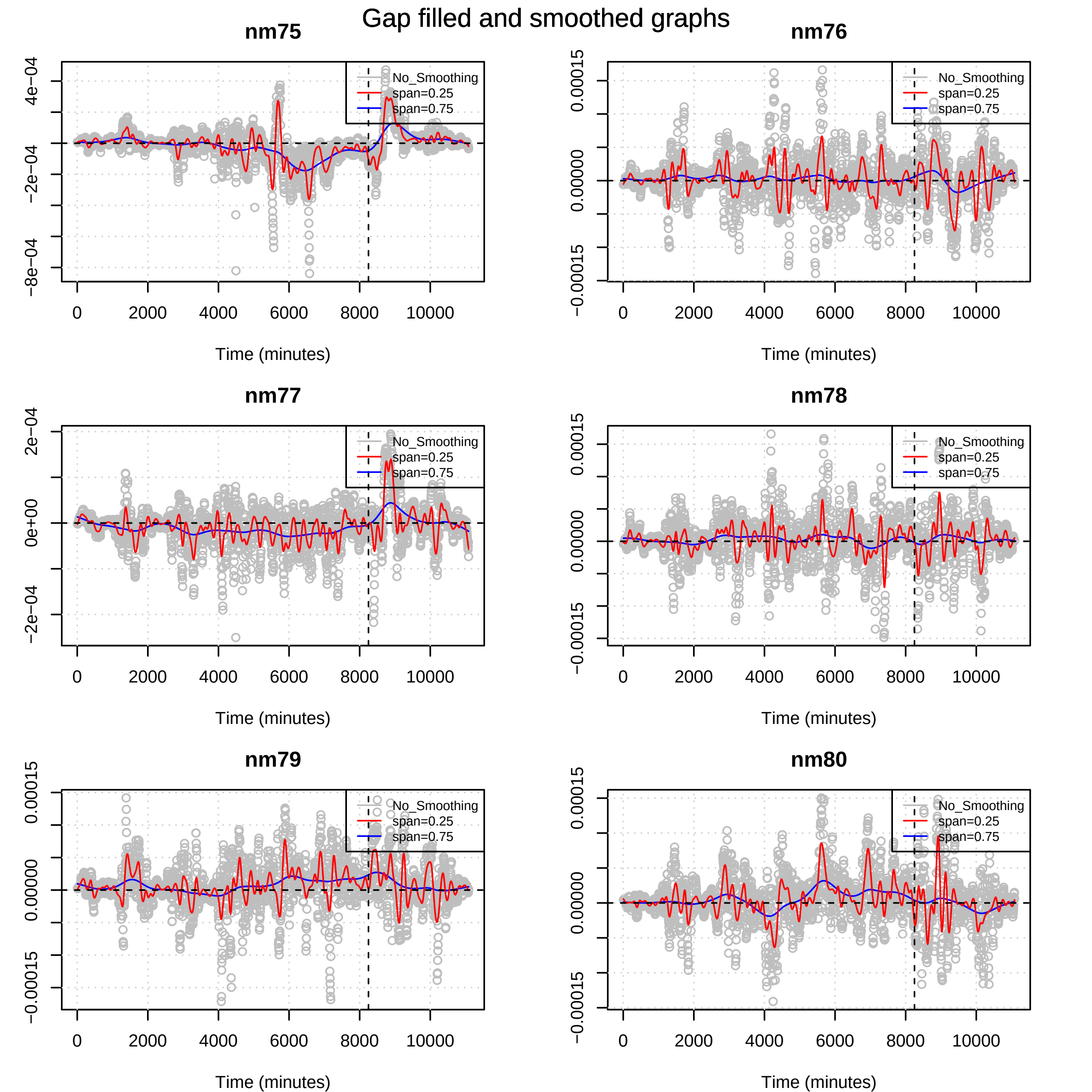

Gap filled and smoothed graphs
nm75
nm76
nm77
nm78
nm79
nm80
No_Smoothing
span=0.25
span=0.75
Time (minutes)
0
2000
4000
6000
8000
10000
4e−04
0e+00
−2e−04
−4e−04
−8e−04
2e−04
0.00015
0.00000
−0.00015

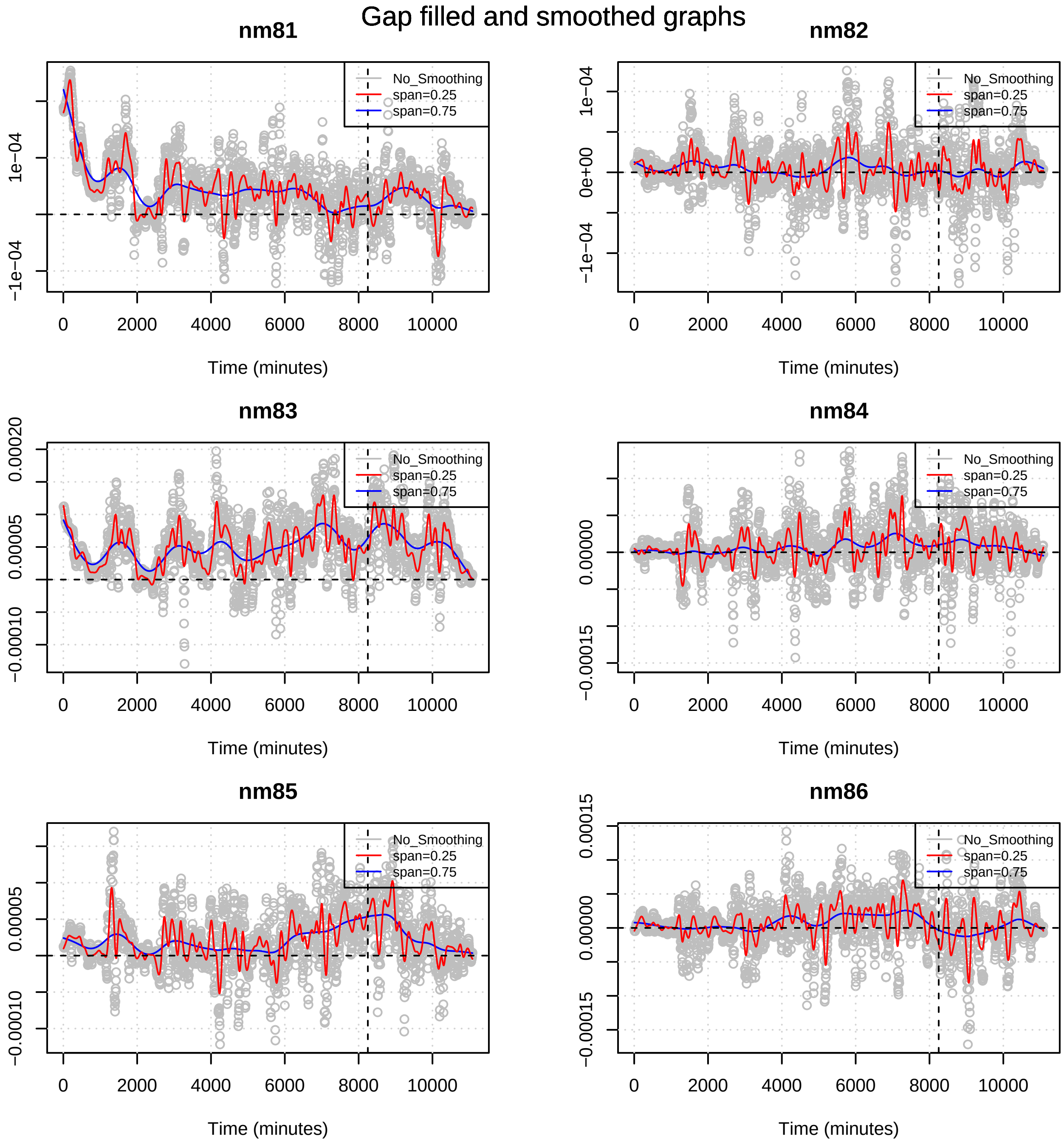

Gap filled and smoothed graphs
nm81
nm82
nm83
nm84
nm85
nm86
No_Smoothing
span=0.25
span=0.75
Time (minutes)
0
2000
4000
6000
8000
10000
1e−04
−1e−04
0e+00
0.00020
0.00005
−0.00010
0.00000
−0.00015
0.00015

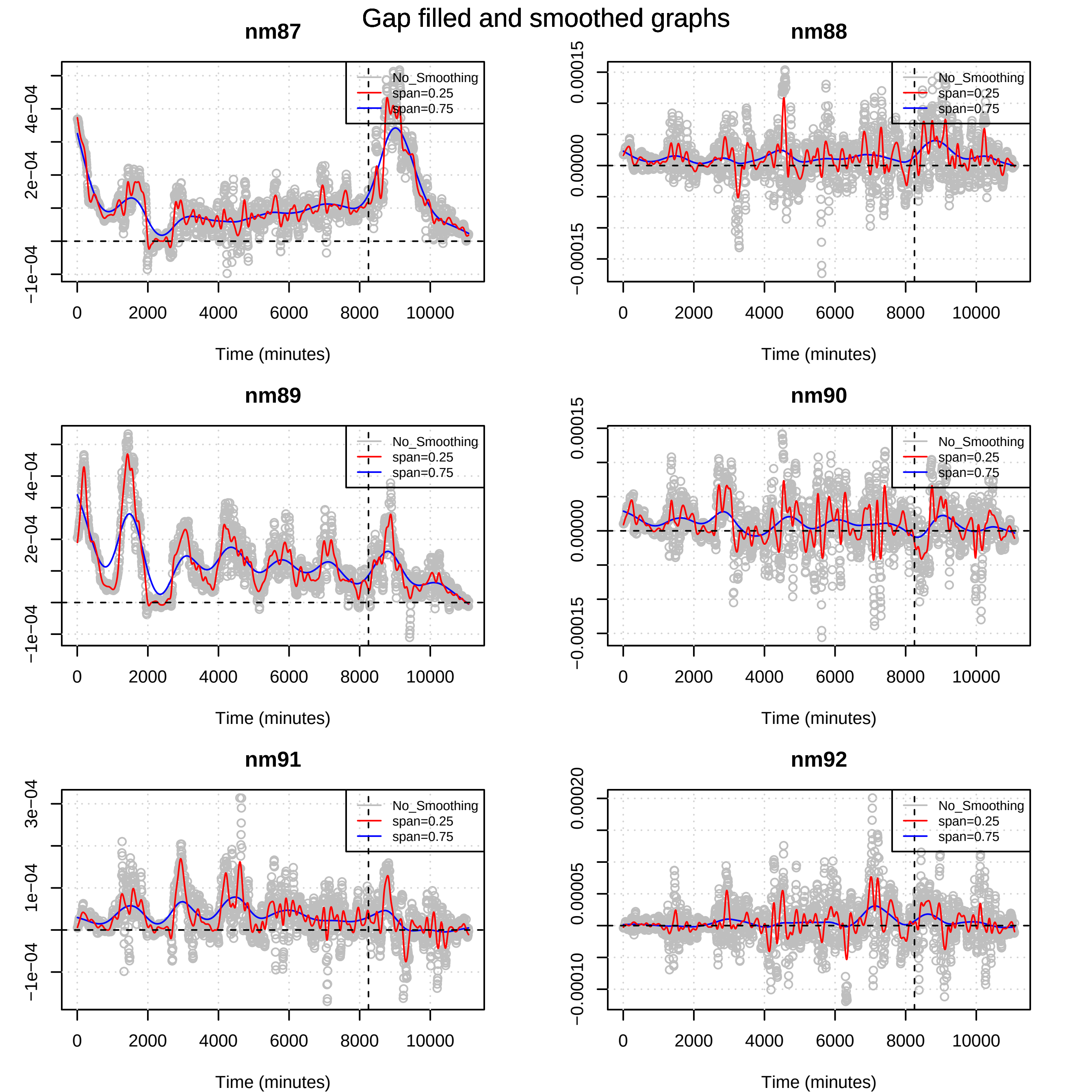
Gap filled and smoothed graphs
nm87
nm88
nm89
nm90
nm91
nm92
No_Smoothing
span=0.25
span=0.75
Time (minutes)
0
2000
4000
6000
8000
10000
−1e−04
2e−04
4e−04
3e−04
1e−04
−0.00015
0.00000
0.00015
−0.00010
0.00005
0.00020

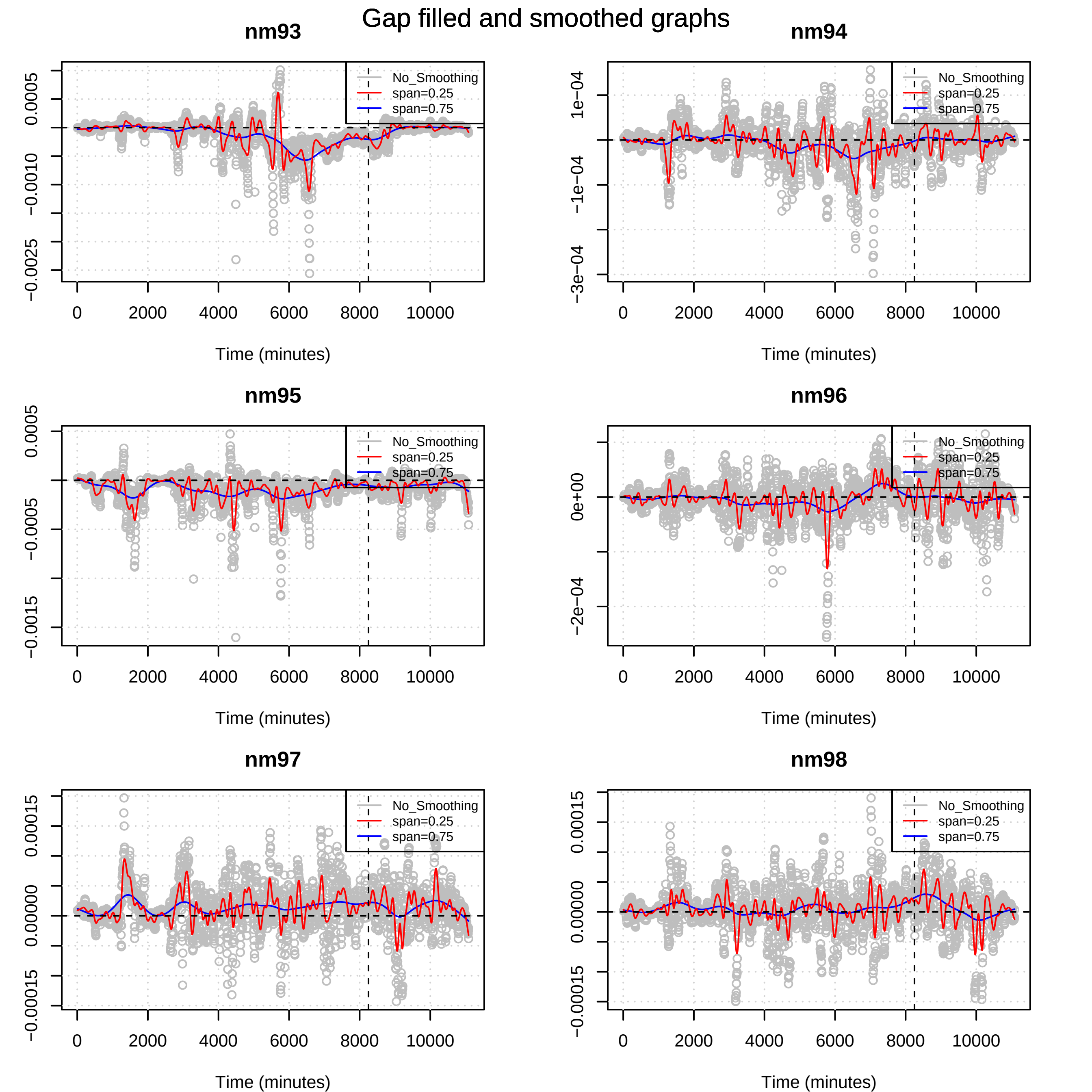

Gap filled and smoothed graphs
nm93
nm94
nm95
nm96
nm97
nm98
No_Smoothing
span=0.25
span=0.75
Time (minutes)
0
2000
4000
6000
8000
10000
0.0005
−0.0005
−0.0010
−0.0015
−0.0025
1e−04
−1e−04
−3e−04
0e+00
−2e−04
0.00015
0.00000
−0.00015

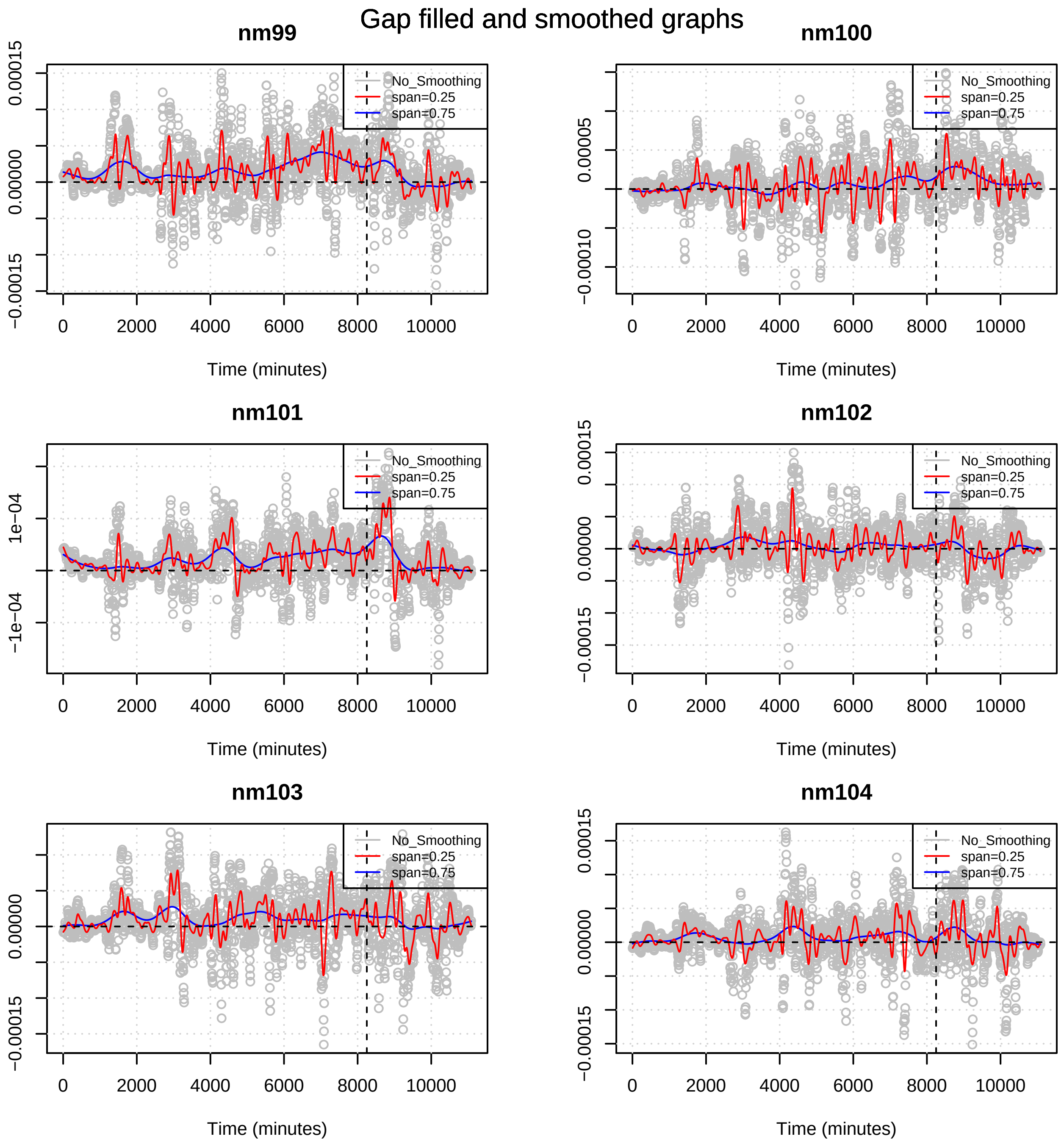
Gap filled and smoothed graphs
nm99
nm100
nm101
nm102
nm103
nm104
No_Smoothing
span=0.25
span=0.75
Time (minutes)
0
2000
4000
6000
8000
10000
0.00015
0.00005
0.00000
−0.00010
−0.00015
1e−04
−1e−04

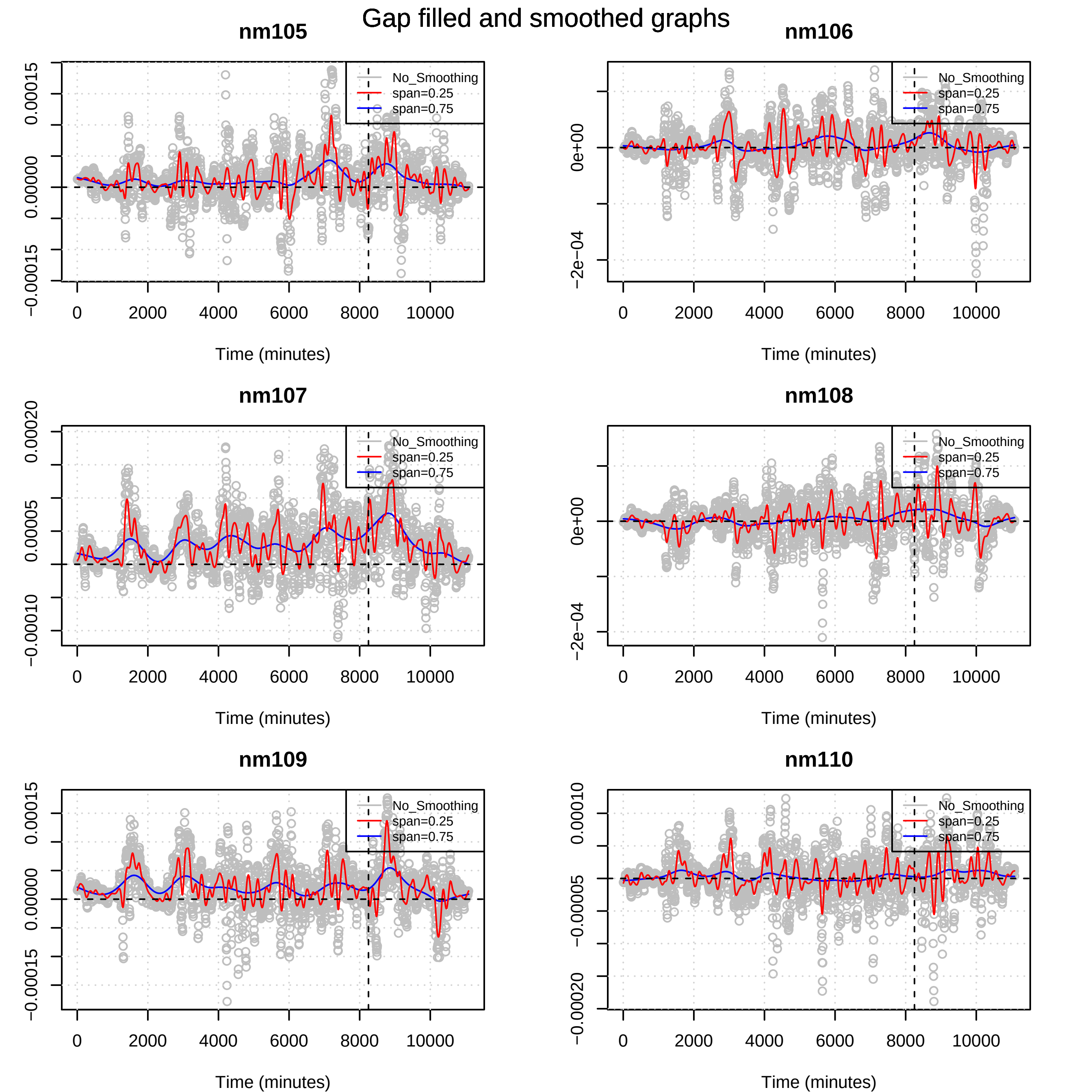
Gap filled and smoothed graphs
nm105
nm106
nm107
nm108
nm109
nm110
No_Smoothing
span=0.25
span=0.75
Time (minutes)
0
2000
4000
6000
8000
10000
−0.00015
0.00000
0.00015
0e+00
−2e−04
−0.00010
0.00005
0.00020
−0.00020
−0.00005
0.00010

### Gap filled and smoothed graphs

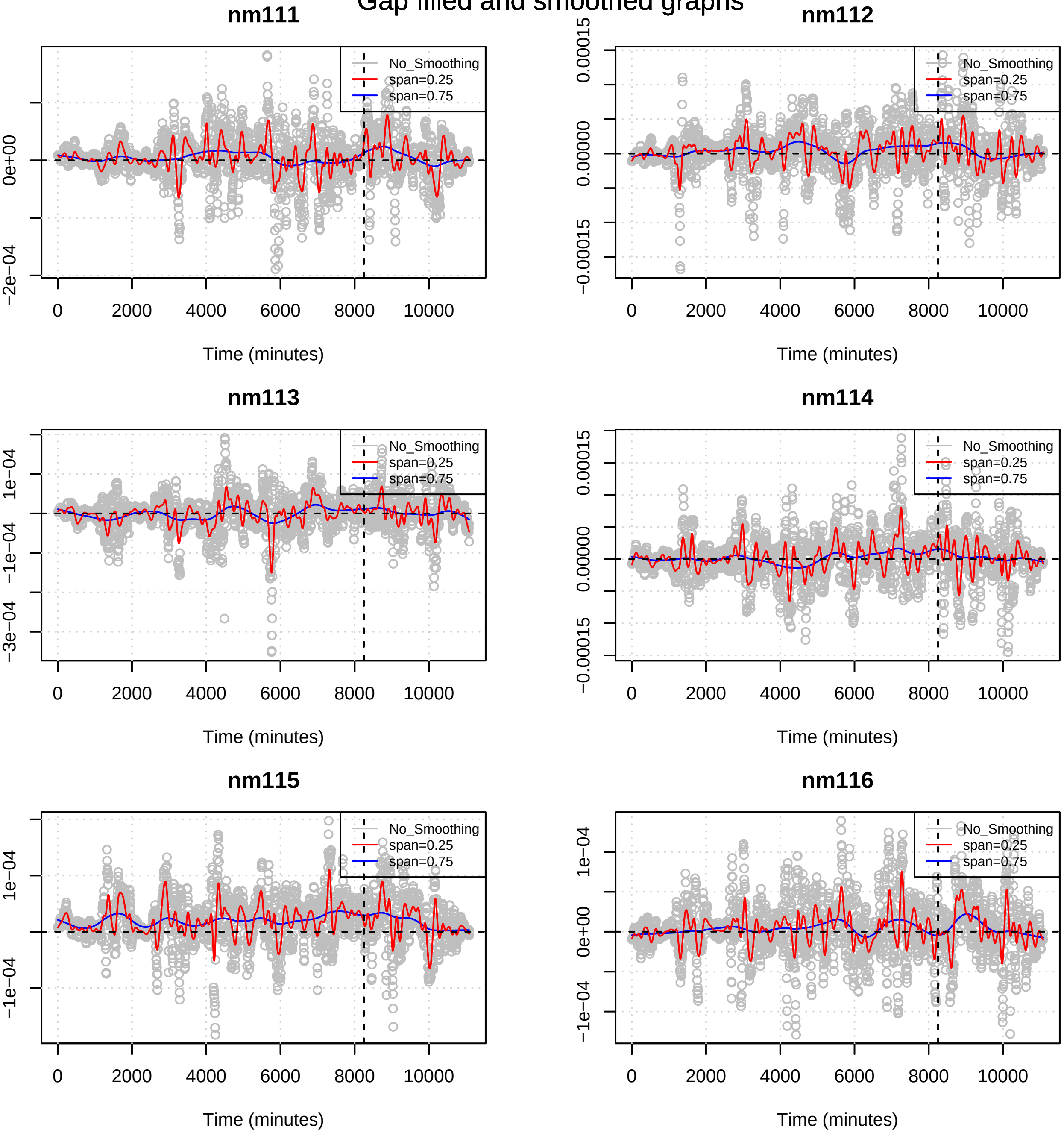

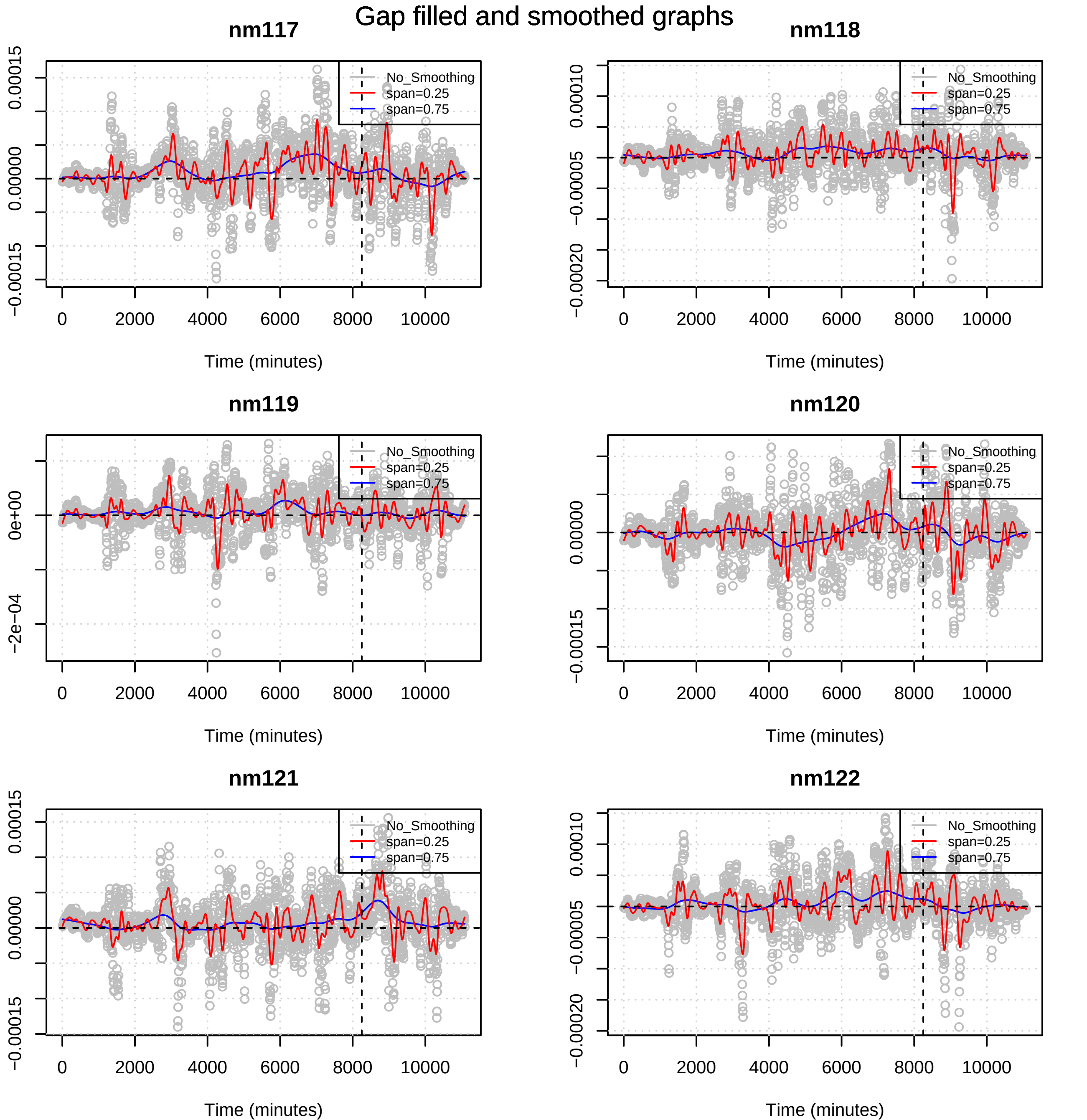
Gap filled and smoothed graphs
nm117
nm118
nm119
nm120
nm121
nm122
No_Smoothing
span=0.25
span=0.75
Time (minutes)
0
2000
4000
6000
8000
10000
−0.00015
0.00000
0.00015
−0.00020
−0.00005
0.00010
−2e−04
0e+00

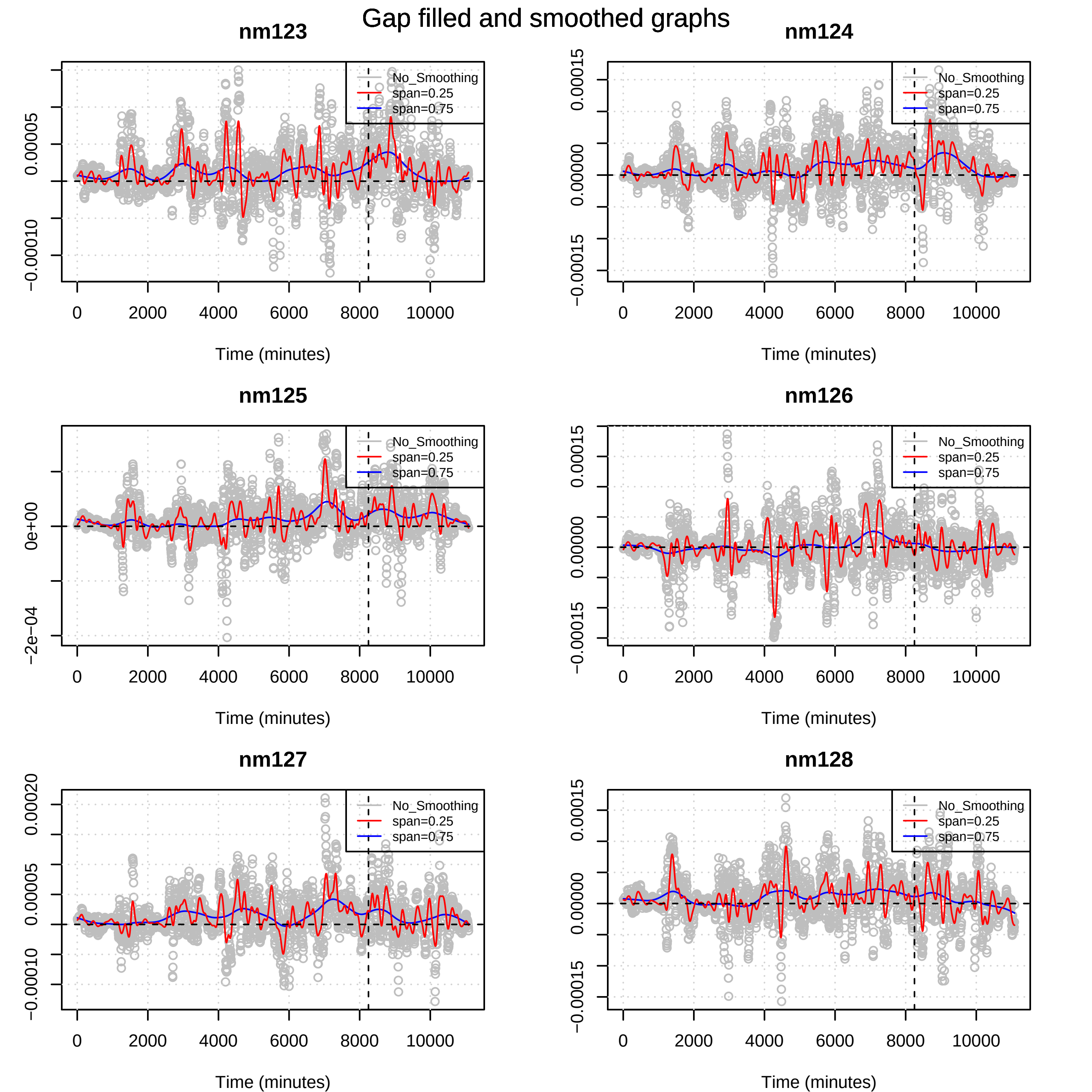

Gap filled and smoothed graphs
nm123
nm124
nm125
nm126
nm127
nm128
No_Smoothing
span=0.25
span=0.75
Time (minutes)
0
2000
4000
6000
8000
10000
-0.00010
0.00005
-0.00015
0.00000
0.00015
-2e-04
0e+00
0.00020

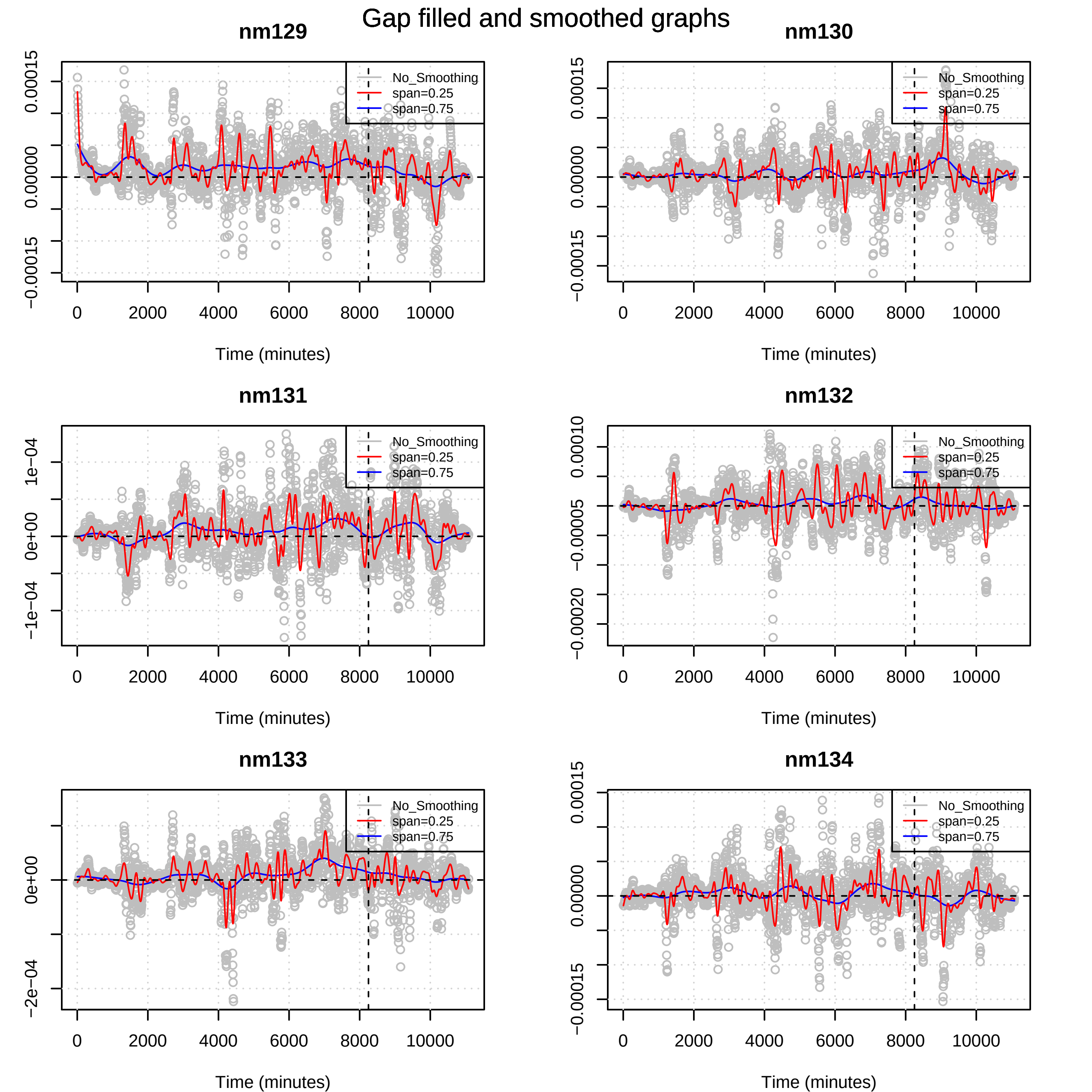

Gap filled and smoothed graphs
nm129
nm130
nm131
nm132
nm133
nm134
No_Smoothing
span=0.25
span=0.75
Time (minutes)
0
2000
4000
6000
8000
10000
−0.00015
0.00000
0.00015
−1e−04
0e+00
1e−04
−0.00020
−0.00005
0.00010
−2e−04

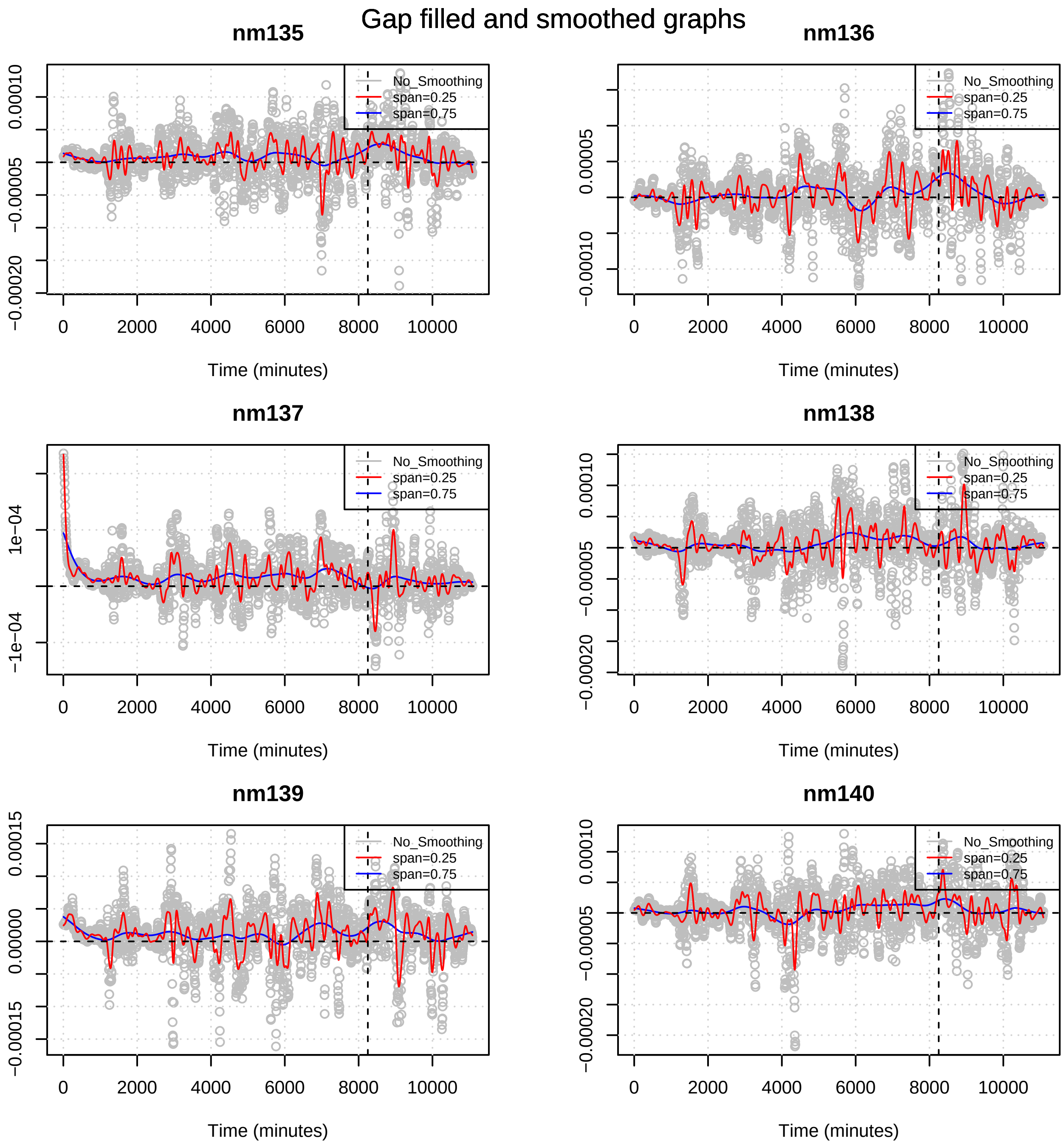
Gap filled and smoothed graphs
nm135
nm136
nm137
nm138
nm139
nm140
No_Smoothing
span=0.25
span=0.75
Time (minutes)
0
2000
4000
6000
8000
10000
0.00010
0.00005
0.00000
−0.00005
−0.00010
−0.00015
−0.00020
0.00015
1e−04
−1e−04

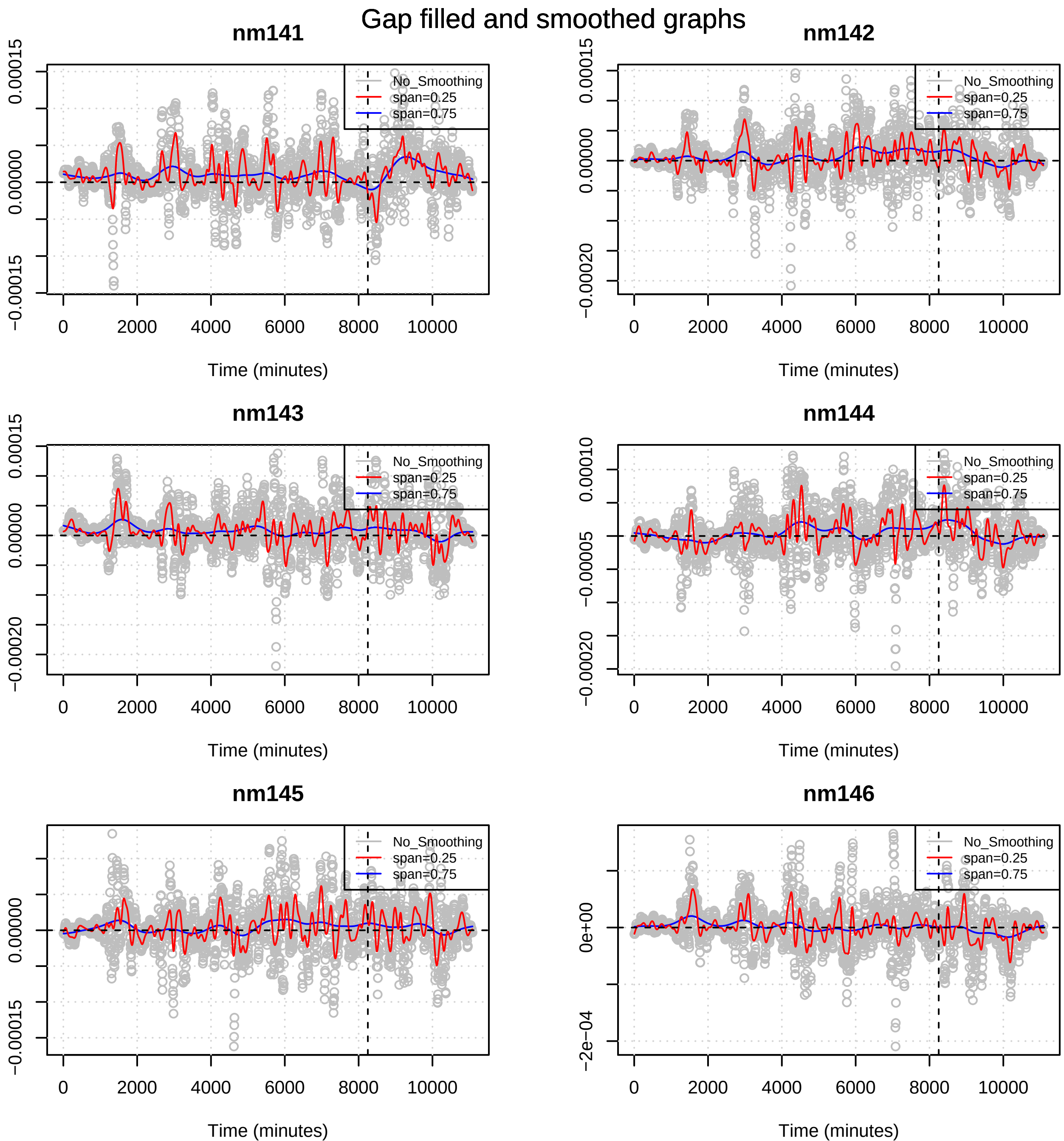

Gap filled and smoothed graphs
nm141
nm142
nm143
nm144
nm145
nm146
No_Smoothing
span=0.25
span=0.75
Time (minutes)
0
2000
4000
6000
8000
10000
0.00015
0.00000
−0.00015
−0.00020
0.00010
−0.00005
0e+00
−2e−04

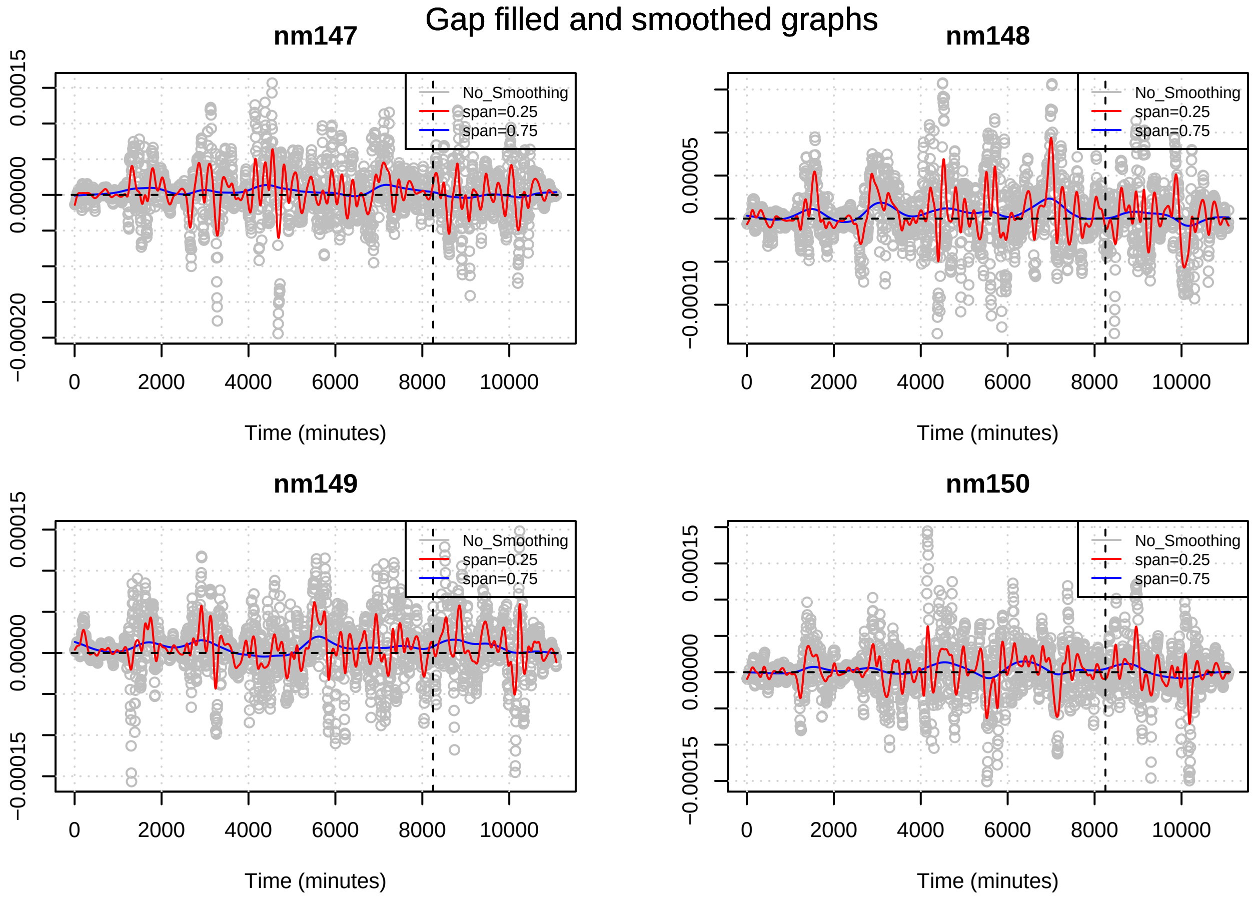

Gap filled and smoothed graphs
nm147
nm148
nm149
nm150
No_Smoothing
span=0.25
span=0.75
Time (minutes)
0
2000
4000
6000
8000
10000
0.00015
0.00000
−0.00020
0.00005
−0.00010
−0.00015
